## Supplementary material for "VolcanoFinder: genomic scans for adaptive introgression": VolcanoFinder: Supporting Information

**Text S1.1** Here, we compare different approximations for the effects of selection on the genealogical distribution at a linked neutral site. As in the main text, $s$ is the (heterozygous) strength of selection acting on the beneficial $B$ allele, $R = rd$ is the rate of recombination between the two sites, and we sample $n = 2$ individuals from a diploid population of size $N$.

**Star-like approximation**

In the main text, the star-like approximation assumes that the stochastic trajectory of the $B$ allele is well approximated by the expected change in allele frequency, *i.e.* logistic growth, from an initial frequency of $1/(2N)$. At a single time point, this average is taken over all possible changes in allele frequency, including a decrease which causes loss of the $B$ allele. In that way, at low frequency, the expected growth is very slow. However, when the allele frequency is very small, its fate – loss or fixation – is largely stochastic. By conditioning on fixation of the $B$ allele, we tend to observe cases where, by chance, the $B$ allele increases in frequency faster than expected. This early stochastic increase can be accounted for by setting the initial frequency to $1/(2N2s)$ [46]. The expected time to fixation is $2\ln(2N2s)/s$, and the probability of escape becomes

$$P_e = 1 - e^{-\frac{R\ln(2N2s)}{s}}. \tag{17}$$

This amounts to re-scaling $\alpha d \to \frac{R\ln(2N2s)}{s}$ in the main text. The same result for $P_e$ was also derived by [135, 136] using a diffusion-approximation approach. The effect of rescaling $\alpha d$ is that the predicted breadth of the sweep increases to fit simulation results more closely. However, this does not account for the fact that $P_{Bb}$ is overestimated while $P_B$ is underestimated by the star-like approximation (Fig. S1.1).

**Dealing with variance in coalescence time**

The fault of the star-like approximation falls in assuming all coalescence occurs at the very beginning of the sweep. In reality, there is variance in the true time to coalescence for the sampled lineages, and late recombination events permit coalescence even to the $b$ background. While this variance in coalescence time can be addressed using a diffusion approach [30], this approximation is valid only for small values of $R/s$. For accurate predictions over the full breadth of the volcano sweep (see Fig. S1.1), we use the approximation in [51], re-derived for the Wright-Fisher (WF) model:

$$P_e = 1 - \frac{s}{R(1-2s)+s}\prod_{j=2}^{M}\left(1 - \frac{R}{js}\right)$$

$$\begin{aligned} P_B &= \frac{s}{R(1-2s)+s}\prod_{j=2}^{M}\left(1 - \frac{2R}{(j+1)s}\right) \\ P_b &= \frac{R(1-2s)}{R(1-2s)+s}\prod_{j=2}^{M}\left(1 - \frac{2R}{(j+1)s}\right) + \sum_{i=2}^{M}\frac{2R}{i(i+1)s}\prod_{j=i+1}^{M}\left(1 - \frac{2R}{(j+1)s}\right) \\ P_{Bb} &= 2\left((1-P_e) - P_B\right) \\ P_{bb} &= 1 - P_B - P_b - P_{Bb} \end{aligned} \tag{18}$$

Here, the establishment of the beneficial allele is modeled as a continuous-time branching process, where the intrinsic birth and death rates are taken as 1/2 (rather than 1 ) to account for drift in the WF (rather than Moran) model [137]. This leads to our term of $R(1-2s)$ rather than $R(1-s)$ in [51]. The subsequent growth of the beneficial allele, conditioned on fixation, is modeled as a pure-birth branching process, or Yule process, which is marked by recombination events to account for the effects of selection on the genealogy at linked neutral sites. If the number of lineages sampled after the sweep is small, their ancestry is well-approximated by the growth of the marked Yule process from the single initial $B$ lineage to $M = 2Ns$ (Moran) or $M = 2N2s$ (WF) lineages. Aside from this factor-of-two difference in $M$, however, the rates of events in the conditioned process are the same in the Moran and WF models.

In Fig. S1.1, we see that the star-like approximation for $(1 - P_e)$, eq. (17), slightly overestimates this but otherwise performs almost as accurately as eq. (18). For comparison, we follow [51] in approximating

$$(1 - P_e) \approx e^{-\frac{R}{s}(\ln(2N2s)+\gamma-2s)} \tag{19}$$

where $\gamma \approx 0.58$ is Euler's gamma. Note that if we ignore $\gamma$ and $s$ terms, we recover the star-like approximation by [46]. This may be interpreted as a better approximation for the time to fixation of the beneficial allele. Indeed $\frac{2R}{s}(\ln(2N2s) + \gamma - 2s)$ closely resembles the approximations in [31] and [137] for the expected time to fixation.

On the other hand, we see in supp. Fig. S1.1 that $P_B$ is underestimated by the star-like approximation, and as a consequence, $P_{Bb} = 2((1 - P_e) - P_B)$ is overestimated. In contrast, eq. (18) estimates $P_B$ very well. We may similarly approximate $P_B$, and by rearrangement and substitution using $1 - P_e$ in eq. (19), we find

$$P_B \approx (1 - P_e)^2 e^{\frac{2R}{s}(1+s)}. \tag{20}$$

The $(1 - P_e)^2$ term corresponds to that of the star-like approximation using $\alpha d = \frac{2R}{s}(\ln(2N2s) + \gamma - 2s)$ as the sweep strength parameter. Importantly, eq. (20) shows us that $P_B$ cannot be accurately approximated using the single sweep parameter $\alpha$. Rather, $P_B$ is $e^{\frac{2R}{s}(1+s)}$ times higher than expected under the star-like approximation.

Let $(1 - P_e)^2$ account for coalescence which occurs at the origin of the sweep and denote $P^* = \frac{(1-P_e)^2}{P_B} = e^{-2R\frac{1+s}{s}}$ the proportion of the $\{B, B\} \to \{B\}$ events which are approximately star-like. Very near the selected site, few if any lineages escape the sweep and most coalescent events will occur very near the origin of the $B$ allele, *i.e.*, as $R \to 0$, $P^* \to 1$ and $P_B \to (1 - P_e)^2$. As $R$ increases, one or both lineages are likely to escape the sweep. Conditioned on sampling lineages with a $\{B, B\} \to \{B\}$ genealogy, coalescence during the sweep occurs only among the subset of non-recombinant $B$ type lineages. $P^*$ decreases to 0 as $R$ grows, and therefore if coalescence occurs, it does so earlier in the sweep than expected under the star-like approximation. However, note that the relative error of the star-like approximation increases with $2R(1 + s)/s$, but $P_B$ decreases more quickly, approximately with $2R\ln(2N2s)/s$. That is, at distances where the error becomes large, $P_B$ is already very small.

**Fig. S1.1 The effect of selection on linked neutral genealogies.** The probability for a lineage not to escape $(1 - P_e)$ and the genealogical distribution for a sample of $n = 2$ as a function of the recombination rate $R$. The solid lines are the approximation of eq. (18). The dashed lines use the star-like approximation with $P_e$ as in 17. The dots represent the average from 1 000 independent simulation runs. **A**. $N = 5\,000$. **B**. $N = 1\,000$. In both panels $s = 0.1$.

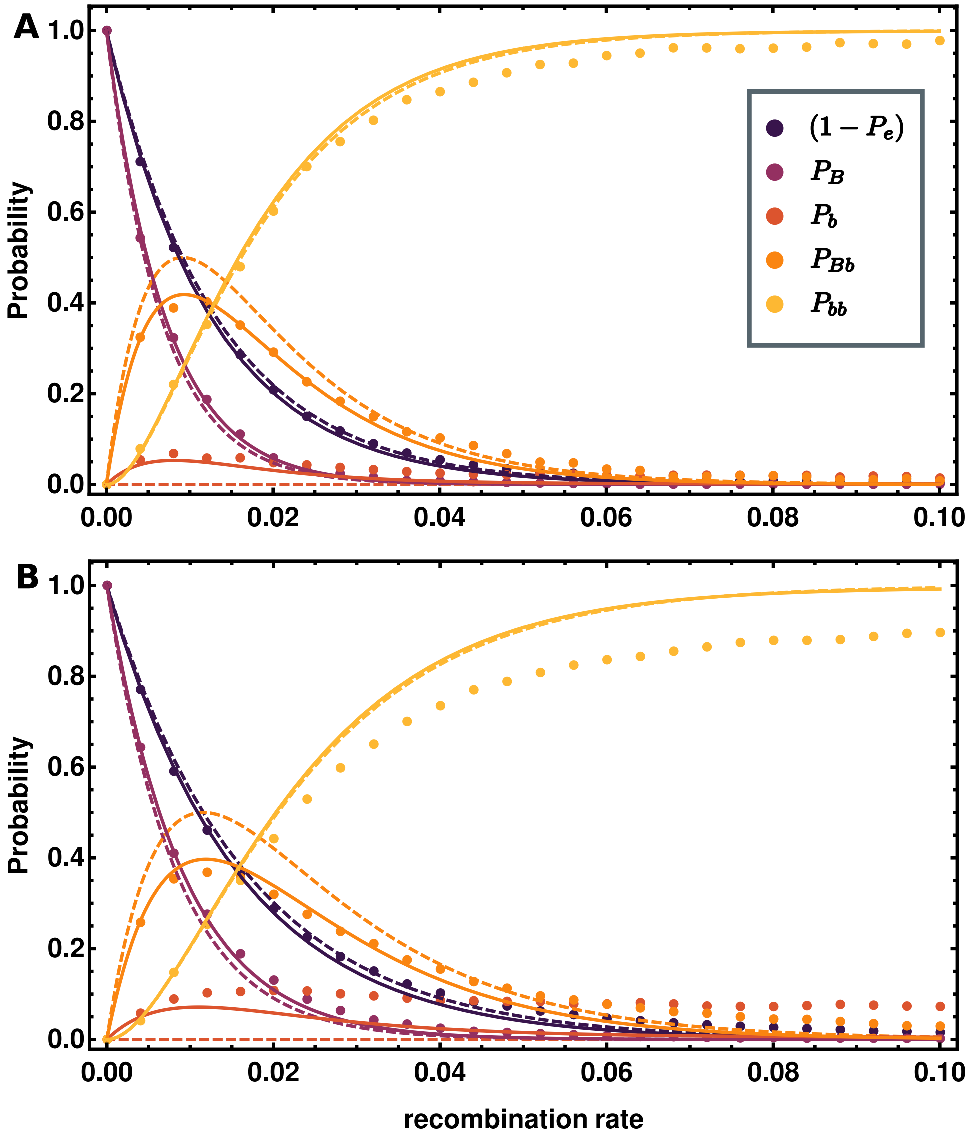

#### Text S1.2 Model 2: accounting for coalescence time within the recipient species.

In a second model we still assume complete lineage sorting, *i.e* all the lineages escaping the introgression sweep coalesce in a single lineage in a recipient species before this lineage coalesce with the single lineage that traced back into the donor species (see Fig. 1), but we no-longer ignore the coalescence time within the recipient species. The $D/2$ factor in the last term in eq. (9) no longer holds and thus needs to be replaced by the probability $\sigma_i(i)$ that a mutation occurred in the ancestral lineage of the $i$ lines that escaped the introgression sweep between the common ancestor and the coalescence event with the lineage that traced back into the donor species. Inspired by eq. (12) we can express the divergences between the recipient and the donor species and between the recipient species and its MRCA with the outgroup species considering the SFS in the subsample of $i$ lineages that escaped the introgression sweep. In the case when fixed differences are polarized we have,

$$\frac{D}{2} = \sigma_i(i) + \sum_{j=1}^{i-1} \frac{j}{i} S_j(i),$$

and

$$D_o = S_i(i) + \sum_{j=1}^{i-1} \frac{j}{i} S_j(i).$$

From these expressions we can isolate $\sigma_i(i) = D/2 - D_o + S_i(i)$ and finally get the probabilities in the altered SFS after the introgression sweep for the polymorphic states $(1 \leqslant i \leqslant n-1)$,

$$S_i'(n|\alpha, d, D) = \left( \sum_{k=i+1}^{n} P_e(k|\alpha, d) S_i(k) \right) + P_e(n-i|\alpha, d) \tfrac{D}{2} + P_e(i|\alpha, d) \left( \tfrac{D}{2} - D_o + S_i(i) \right). \tag{21}$$

Eq. (21) is only valid if $D/2 - D_o + S_i(i) \geqslant 0$ for all $i \in \{1, \ldots, n\}$. A necessary and sufficient condition is $\frac{D}{2} \geqslant D_o - S_n(n)$. Using eq. (12) leads to $\frac{D}{2} \geqslant \frac{n-1}{n} \hat{\theta}_L$, where $\hat{\theta}_L = \frac{1}{n-1} \sum_{i=1}^{n-1} i S_i(n)$ is an unbiased estimator of $\theta$ defined in [52, eq.(8)] and computed from the whole genomic background.

Similarly, the $D_o$ factor in the last term of eq. (11) can easily be replaced by the probability $S_n(n)$ that a mutation occurs on the ancestral lineage of $n$ lineages that have escaped the introgression sweep before coalescence occurs with the outgroup:

$$S_n'(n|\alpha, d, D) = \left( \left( D_o - \tfrac{D}{2} \right) \sum_{k=1}^{n-1} P_e(k|\alpha, d) \right) + D_o P_e(0|\alpha, d) + S_n(n) P_e(n|\alpha, d). \tag{22}$$

If fixed differences are not polarized, then eqs. (21) and (22) still hold when substituting the divergence between the recipient species and its MRCA with the ougroup species $D_o$ with the full divergence between the recipient and the outgroup species $D_o'$. Once again the probabilities in eqs. (21) and (22) are linearly dependent of the mutation parameter $\theta = 4N\mu$, and this dependency disappears in conditional probabilities obtained from eqs. (10) and (13).

### Text S1.3 Comparison of Models 1 and 2

In Fig. 4 of the main text, we saw that Models 1 and 2 yield similar predictions for the SFS after the selective sweep when $D \gg \theta$. That is, when the divergence is sufficiently large so that ancestral variation is no longer segregating in the populations, $T_{coal,2} \ll T_d$, the pairwise diversity $D$ well-approximates the contribution of fixed derived mutations from the recipient population. Here, we look at the difference between the two models in approximating the expected heterozygosity after the sweep.

For a sample of $n = 2$ two lineages taken directly after the sweep, the expected heterozygosity may be approximated as in eq. (3) of the main text. In Fig. S1.2, we show the star-like approximation in grey, the more-accurate approximation of [51] in black, and simulation results as black dots. For a sample of $n > 2$, we use the un-normalized $S'_i(n)$ from either Model 1 (eq. 9) or 2 (eq. 21) to determine the effects of the sweep. By substituting the $S'_i(n)$ into eq. 6, the expected heterozygosity is given by $S'_1(2)$. We show the predictions of Model 1 (red, dashed) and Model 2 (blue, dashed) in Fig. S1.2. We see that Model 2 exactly matches the predictions of the star-like approximation and that Model 1 is even more biased to over-estimates the increase in genetic diversity.

### Text S1.4 The performance of `VolcanoFinder` and `SweepFinder`

Here we take a closer look at the ability of both `SweepFinder` and `VolcanoFinder` to detect an adaptive introgression sweep. 200 successful introgression sweeps centered in a 500 kb locus were simulated under strong selection, $2Ns = 1\,000$ ($N = 5\,000$, $s = 0.1$), a scaled mutation parameter $\theta = 0.002$ (mutation rate $\mu = 10^{-7}$), and significant divergence of the donor population $D = 0.026 = 13\theta$. The per-site recombination rate $r = 5 \times 10^{-7}$, thus the sweep parameter $\alpha = r\ln(2N)/s = 4.6 \times 10^{-5}$. The data consists of $n = 10$ lineages sampled from the recipient species. It is polarized to an outgroup with pairwise divergence $D_o = 0.05$ and includes fixed differences. As shown in the power analysis of the main text, both sweep scan methods have high power to detect introgression sweeps with these parameters.

Single iterations of the adaptive introgression process may produce the expected volcano pattern, but when early recombination events occur, the signal is concatenated. In order to compare data to theory, the 200 iterations were combined into a single data-rich locus, preserving the unique identifier and correct positions of the mutations within the simulated genomic region The result is an "average" data set representative of the expected volcano sweep pattern which we scan for selection using `SweepFinder` and `VolcanoFinder`. Note that here, the LR values are inflated because they are calculated from the combination of 200 iterations of data. The LR values of a single iteration are much smaller.

`VolcanoFinder` scans were run over a range of potential divergence values $D = 0.010$, 0.015, 0.020, 0.026, 0.030, 0.035, 0.040, and 0.045 (*i.e.* $D/\theta = 5$, 7.5, 10, 13, 15, 17.5, 20, 22.5) including the true value used in simulations $D_{sim} = 0.026$. While the true value $D = D_{sim}$ results in a very high LR value of 507297, `VolcanoFinder` finds that an introgression sweep with $D = 0.020$ fits the average data slightly better, with LR value 598280. As shown in the previous section, model 1 consistently over-estimates the contribution of divergence before the sweep to diversity after the sweep. In combination, the star-like approximation for the effect of selection on linked neutral loci systematically over-estimates the height of the diversity peaks. Together, this indicates that a lower $D$ value yields a better-fitting model. Indeed, supp. Fig. S1.3 shows that at the distance $\alpha d = 1/2$ where the volcano peak is close to the maximum height, the model predictions fit the data better using $D = 0.02$. However, for both $D$ values, `VolcanoFinder` finds an optimum sweep parameter

$\hat{\alpha} \approx 3.4 \times 10^{-5}$, close to the true value of $4.6 \times 10^{-5}$, ($\frac{\hat{\alpha}}{\alpha} \approx 0.74$).

`SweepFinder` is also able to detect the introgression sweep but is less sensitive to the signal, producing a LR value of 2672, nearly 200 times smaller than that reported by `VolcanoFinder`. While `VolcanoFinder` uses information from the full breadth of the introgression sweep, the `SweepFinder` model cannot account for the influx of foreign variation and is sensitive only to the features of the narrow diversity valley near the introgression sweep center.

In Fig. S1.4, we compare the optimum model that `SweepFinder` fits to the average data set to that of `VolcanoFinder` with $D = 0.026$, discussed above and shown in Fig. S1.3 (right column). In the top panel, we see that `SweepFinder` detects only a very small sweep valley that approximately matches the valley of the volcano sweep. In contrast to `VolcanoFinder`, the optimum sweep strength found by this method is an order of magnitude smaller than the true value used in simulations ($\frac{\hat{\alpha}}{\alpha} \approx 6.4$).

As expected, the weak sweep parameter chosen by `SweepFinder` as the optimum allows it to approximately fit the SFS very near the sweep center (top two panels, classic sweep model $\hat{\alpha}d = 0.01$ or 0.1). At greater distances, `SweepFinder` cannot predict the effect of introgression on the SFS and matches the data poorly. Due to the weak sweep strength, the corresponding regions in the simulated data are in-reality an order of magnitude closer to the sweep center when distance is scaled by the true strength of the selective sweep. This has conflicting effects on the power of `SweepFinder` to detect adaptive introgression sweeps.

At distances $\alpha d > 10$ from the sweep center, a selective sweep has little effect on neutral genealogies, and this provides a limit to how much of the data is informative for selective sweep scans. With a much weaker selection strength parameter, `SweepFinder` assesses only a fraction of the information that is accessible to the `VolcanoFinder` method, explaining in-part the much-lower LR value.

However, by finding a weak optimum strength parameter, `SweepFinder` avoids looking at regions of the introgression sweep in which it performs poorly *relative to the background SFS used as a null hypothesis*. In the main text, we saw that at distances greater than or equal to $\alpha d = 1$, the classic sweep model predicts a near-return to the background SFS. At greater distances, sites are no longer informative due to the similarity in the null and alternative hypotheses of the likelihood ratio statistic.

**Fig. S1.2 Pairwise diversity after the selective sweep.** Predictions from a sample of two lineages are in gray. Model 1 predictions are in red. Model 2 predictions are in blue. Our original model predictions are in black. Average of simulated data points ±3 standard error are shown in black. In the upper data set $D = 0.026$, and in the lower data set $D = 0.014$. In both, the remaining parameters are $\theta = 0.002$, $N = 5\,000$, $s = 0.1$, and $n = 50$.

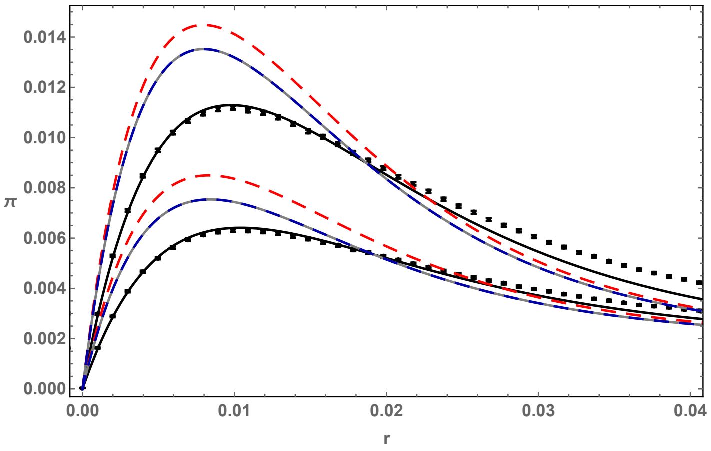

**Fig. S1.3** `VolcanoFinder` **optimum model: choice of** $D$ Results from the `VolcanoFinder` scan of the average data set described in the supp. Text S1.4. The left column shows the optimum sweep model (inferred strength parameter $\hat{\alpha} = 3.4 \times 10^{-5}$) given the true $D = D_{sim} = 0.026$ used in the simulation. The right column shows the best-fitting model found by the method with inferred parameters $\hat{D} = 0.02$ and $\hat{\alpha} = 3.4 \times 10^{-5}$. The top panels show the average heterozygosity along the sweep region with distance scaled by the true sweep strength $\alpha d$ (gray) as well as the expected diversity predicted under the given model (blue dashed). The remaining rows show the theoretical SFS of the the optimum model (light gray) at increasing distances $\hat{\alpha} d = 0.01, 0.1, 0.5, 1, 2, 3, 8$ from the sweep center and compares this to the observed SFS in a 100-bp window centered at that position averaged over 50 simulations (dark gray). The label on each panel lists the chosen value of scaled distance $\hat{\alpha} d$ from the optimum model as well as the corresponding *true* value of $\alpha d$ determined by the simulation parameters.

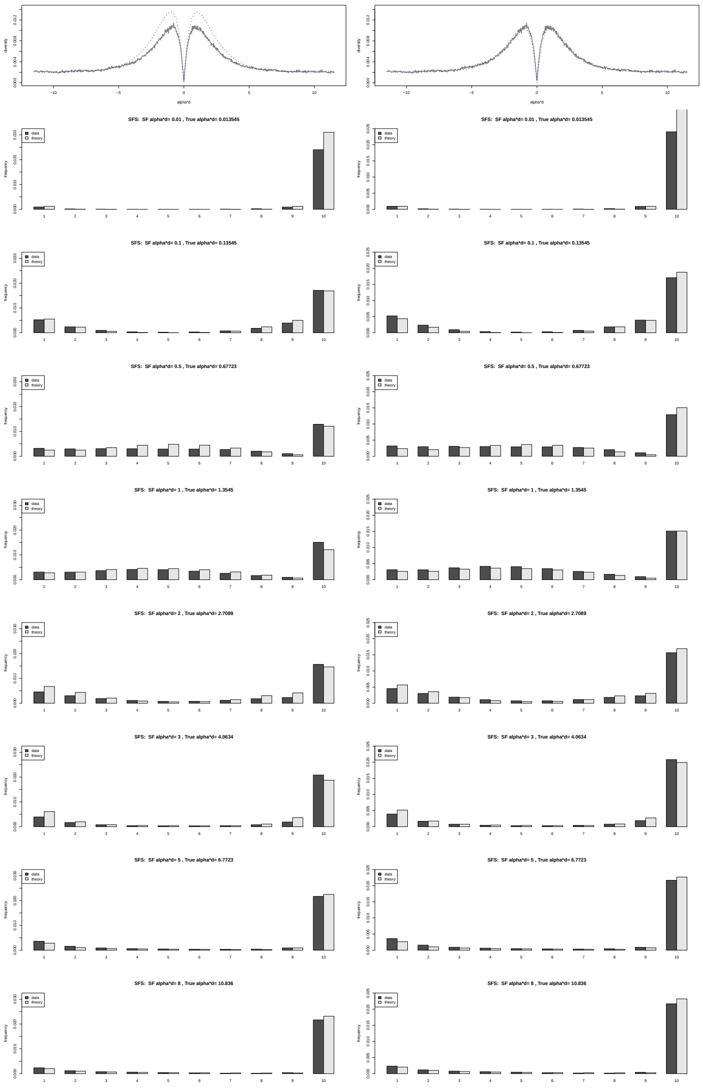

**Fig. S1.4** `SweepFinder` **detects the introgression sweep valley.** The left column shows the best-fitting model for `SweepFinder`. The `VolcanoFinder` results in the right column as well as the description of the panels are the same as in Fig. S1.3.

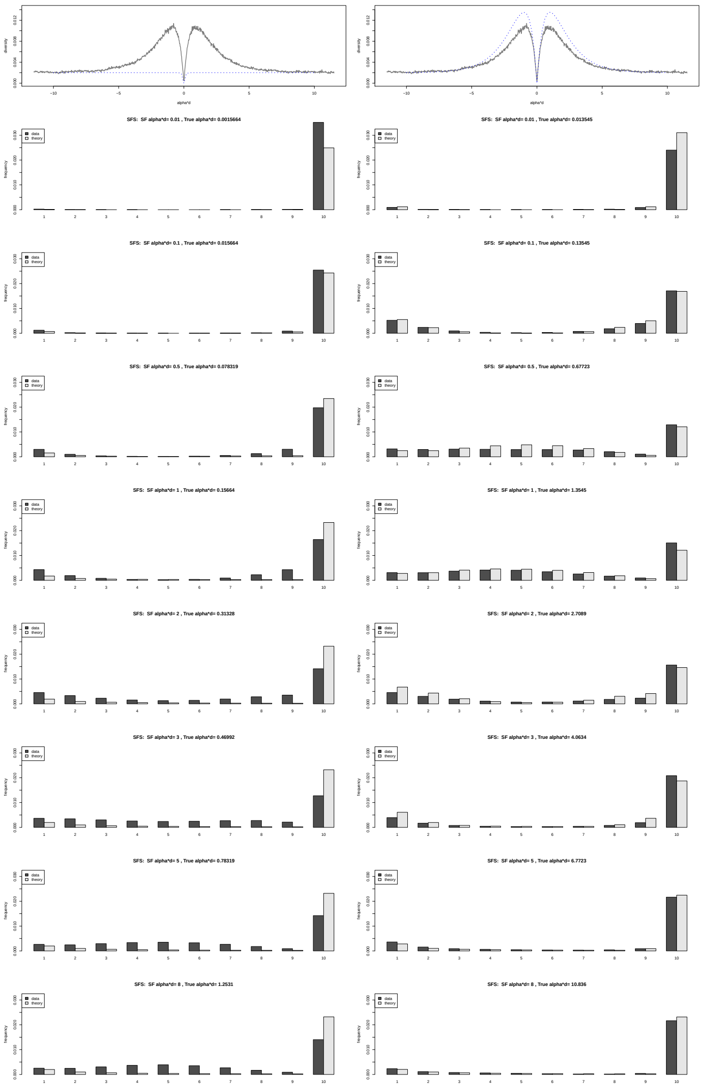

### Text S2.1 Adjusting the migration and duration for the introgression sweep.

We use the model in Fig. S2.1, where introgression from a donor species occurs as a short and sudden migration episode from the donor to the recipient species. In order to simulate the sweep of an introgressed selected allele, we adjust the migration parameter $m$ such that the expected frequency $p_0$ of the introgressed selected allele just after the migration episode leads to the fixation of the selected allele with a high probability $\pi_{\text{fix}} = 0.95$.

**Migration rate**. Under a diploid additive model the probability of ultimate fixation of a beneficial allele with initial frequency $p_0$ is given by eq. (5.47) in [138]

$$\pi_{\text{fix}} = \frac{1 - e^{-2Nsp_0}}{1 - e^{-2Ns}}. \tag{23}$$

Isolating $p_0$ in eq. (23) leads to

$$p_0 = -\frac{1}{2Ns} \ln\left(1 - \left(1 - e^{-2Ns}\right) \pi_{\text{fix}}\right). \tag{24}$$

In a discrete generation model, the backward migration rate $m$ is defined as the probability for a lineage in the recipient population to come from the donor population at the previous generation, we thus have $p_0 = m$ and adjust the migration rate to obtain a desired fixation probability for the selected allele. In a continuous coalescent model where time is scaled in units of $4N$ generations, a comparible result is obtained by seting the migration rate to $m = p_0/\Delta t_{\text{mig}}$ during a time interval $\Delta t_{\text{mig}}$.

**Duration of the sweep phase**. We assume weak selection such that $\bar{w} \approx 1$ and the dynamics of the frequency of the beneficial allele is well described by the logistic model [138, eq. (1.27)]. If the time $t$ is scaled in units of $4N$ generations, we have

$$\frac{\mathrm{d}p}{\mathrm{d}t} = 4Nsp(1-p) \Rightarrow t = \frac{1}{4Ns}\left(\ln p - \ln(1-p)\right) + C, \tag{25}$$

where $C$ is an integration constant. The duration of the sweep phase $\Delta t_{\text{sweep}}$ can be computed from the frequencies of the beneficial mutation at the introgression time $p_0$ and at the end of the sweep $p = 1 - 1/(2N)$:

$$\Delta t_{\text{sweep}} = \frac{1}{4Ns}\left(\ln\left(1 - \frac{1}{2N}\right) - \ln\frac{1}{2N} - \ln p_0 + \ln(1-p_0)\right). \tag{26}$$

Including drift at the end of the sweep would substantially shorten the duration of the sweep, and using eq. (26) enables that the selected went to fixation with a high probability.

### Text S2.2 Coalescent simulations.

The coalescent simulations were conducted with the `R` package `coala` [72] as a frontend to the coalescent simulator `msms` [69].

#### Coalescent simulations involving introgression sweeps.

We used the model described in supp. Fig. S2.1. We simulated coalescent trees for $n = 40$ lineages sampled from the recipient species and one from the outgroup species. The simulated region comprises $2 \times 10^5$ nucleotides, the recombination rate was set to $\rho = 5 \times 10^{-7}$ events per generation per nucleotide and the mutation rate was set to $\mu = 1.25 \times 10^{-7}$ events per generation per nucleotide following previous studies [49]. The speciation time between the outgroup and the ancestor of the donor and recipient species was $T_{sp} = 10$ units of $4N$ generations (8 Mya with $N = 10^4$ and a 20 years generation time). The divergence time between the donor and recipient species was $T_d \in \{1,\ 2.5,\ 4,\ 5.5\}$ ($D/\theta \in \{3,\ 6,\ 9,\ 12\}$, equivalent to 0.8, 2, 3.2, and 4.4 Mya, respectively). Taking into account an average expected time of $2N$ generations for the coalescence of a pair of lineages coming for the donor and recipient species, these values of $T_d$ lead to an average probability for a nucleotide to differ between the donor and the recipient species (average divergence) of $D \in \{0.015,\ 0.03,\ 0.045,\ 0.06\}$. The population sizes of the recipient species, the ancestor of the recipient and donor species, as well as the outgroup species were set to $N = 10^4$ individuals. To mimic a sweep from a single lineage introgressed from the donor species (hard introgression sweep), the population size of the donor species was reduced to a single individual ($N_d = 1$) at the time of the split such that all lineages that trace back into the donor species coalesce almost instantaneously. We also relaxed this assumption, restraining this bottleneck to a tiny time interval (20 generations) after the split from the recipient species and thus allowing the donor species to recover polymorphism before the introgression event, potentially allowing sweeps from different lineages in the recipient species (soft introgression sweeps). Because we do not model an initial selective sweep in the donor population, the nucleotide diversity in the donor population is only affected by the strong bottleneck and is not locally reduced around the selected site.

Introgression from the donor into the recipient species occurs at time $T_i = T_s + \Delta t_{\text{sweep}} + \Delta t_{\text{mig}}$: migration is allowed from the donor species into the recipient species for a short time interval $\Delta t_{\text{mig}}$. The migration rate $m$ and duration of the migration interval $\Delta t_{\text{mig}}$ are chosen such that the expected frequency of the selected allele in the recipient species at the end of the migration interval leads to the fixation of the selected allele with a high probability $\pi_{\text{fix}}$ as described in Text S2.1. Two selection coefficients were used $2Ns \in \{100,\ 1000\}$ and $\pi_{\text{fix}} = 0.95$ was achieved by using an identical migration rate in both cases ($m \approx 0.003$) and letting the duration of the migration event last a single generation ($\Delta t_{\text{mig}} = 1/(4N)$ for $2Ns = 1\,000$) or 10 generations ($\Delta t_{\text{mig}} = 10/(4N)$ for $2Ns = 100$). A logistic model for the dynamics of the selected allele as in eq. (25) eventually leads to fixation after some time given by eq. (26). We assume that sampling was done at time $T_s \in \{0,\ 0.1,\ 0.25,\ 0.5\}$ (equivalent to 0, 80, 200, and 400 kya respectively) after the fixation event.

*Total number of replicates.* Combining four values for the divergence time $T_d \in \{1,\ 2.5,\ 4,\ 5.5\}$, four values for the time since the selective sweep $T_s \in \{0,\ 0.1,\ 0.25,\ 0.5\}$ and two values for the selection coefficient $2Ns \in \{100,\ 1\,000\}$ for each hard and soft introgression sweeps leads to 64 parameter sets. For each parameter set, we ran 1 000 coalescent simulations involving selection and 10 000 neutral coalescent simulations with the same admixture level (same migration rate at the same time point). In addition, a neutral non-admixed reference was obtained from another

10 000 coalescent simulations. In total we thus performed 714 000 (650 000 neutral and 64 000 non-neutral) coalescent simulations.

**Coalescent simulations involving balancing selection.**

We used three different demographic models (Fig. S2.2) inspired by the models that were formerly used to investigate the statistical power of `BALLET` [48]. The speciation time between the outgroup and the ingroup species was $T_{sp} = 10$ units of $4N$ generations (8 Mya with $N = 10\,000$ and a 20 years generation time). The simulated sequence comprised $2 \times 10^5$ nucleotides. Balancing selection involved a selected locus with two alleles (A and a) in the middle of the sequence. We assumed overdominance with the following fitnesses: $w_{aa} = 1$, $w_{Aa} = 1 + hs$ and $w_{AA} = 1 + s$ with $h = 100$ and $s = 0.01$. Balancing selection started at different times $T_s \in \{1.25, 5, 8.75, 12.5, 16.25, 20\}$ (equivalent to 1, 4, 7, 10, 13, and 16 Mya, respectively). We assume the selected allele $A$ reaches the equilibrium frequency $h/(2h - 1)$ as soon as selection starts. For the population growth model, population expanded from $N = 10\,000$ to $N_g = 2N$ at time $T_g = 0.06$ (48 kya). For the bottleneck model, population size was reduced from $N = 10\,000$ to $N_b = 0.055N$ at time $T_b = 0.0375$ (30 kya) before returning to its initial size at time $T_e = 0.0275$ (22 kya).

*Total number of replicates.* Combining three demographic models and six values for $T_s$ leads to 18 parameter sets. We ran 1 000 coalescent simulations involving selection for each parameter set, and 10 000 neutral coalescent simulations for each demographic model. In addition to these simulations, a single simulation of a $2 \times 10^7$ nucleotides sequence was performed to generate a genomic background reference for each demographic model. In total we thus performed 48 003 (30 000 neutral, 18 000 non-neutral and 3 neutral background references) coalescent simulations.

**Fig. S2.1 Introgression model.** The model comprizes three species. The ancestor of the donor and recipient species diverged from an outgroup species at time $T_{sp}$ in the past. The donor and recipient species diverged at time $T_d$ in the past. Immediately after this speciation event, selection starts (diploid additive model with selection coefficient $s$) in the donor and recipient species, but the beneficial allele is only present in the donor species, where it is assumed to have already reached fixation. The donor species is bottlenecked to a population size of $N' = N_d$ individuals ($N_d$ is assumed to be very small so that coalescence of lineages in the donor species is immediate). The bottleneck may last until present (enforcing a hard introgression sweep, vertical dashed line) or only occur for a very short period after which the donor population size is set to $N' = N$, allowing the donor population to recover some polymorphism before the introgression occurs (soft introgression sweep). At time $T_i$ in the past, migration occurs for a small amount of time (one to ten generations) from the donor to the recipient species. At this time point the beneficial allele is introgressed in the recipient population. The migration rate $m$ is set such that the fixation probability of the beneficial allele in the recipient population is 0.95 given its expected initial frequency in the recipient population at time $T_i$. The selected allele reaches fixation in the recipient species at time $T_s$.

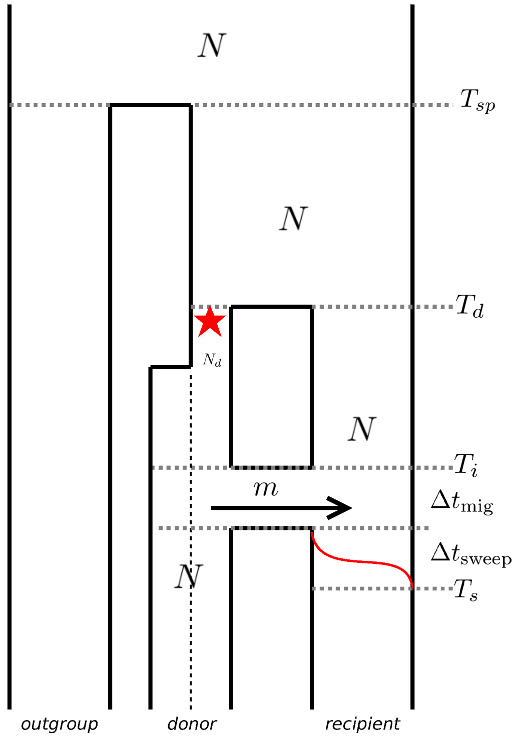

**Fig. S2.2 Balancing selection models.** The model comprizes two species, the outgroup and the ingroup species that diverged at time $T_{sp}$ in the past, when the ancestor species (size $N_a$) splitted into two species of equal sizes $N = N_a$. Balancing selection starts at time $T_s$ either in the ingroup species ($T_s < T_{sp}$) or in the ancestor species ($T_{sp} < T_s$). **A** Constant population size model. **B** Population growth model: the size of the ingroup species expanded to $N_g$ at time $T_g$. **C** Bottleneck model: the size of the ingroup species was reduced to $N_b$ between times $T_b$ and $T_e$.

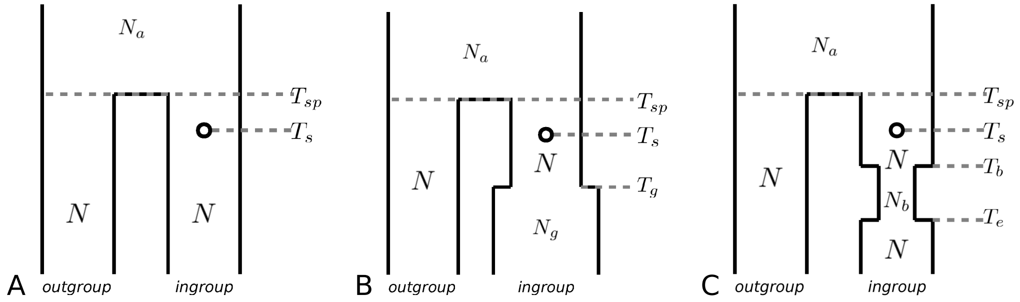

### Text S2.3 Genome scans and power analysis.

#### Genome scans.

*Genome scans for introgression sweeps.* We compared the performance of three model-based composite likelihood methods: `BALLET` [48], `SweepFinder2` [49] and `VolcanoFinder`. For each method, two cases were considered for the reference genomic background. We considered either a non-admixed reference inferred from 10 000 neutral coalescent simulations without introgression or an admixed reference inferred from 10 000 neutral coalescent simulations involving the same level of admixture as the non-neutral coalescent (same migration rate at the same time point) as described in Text S2.2 and Fig. S2.1.
`BALLET`: the $T2$ test of `BALLET` was used with a sliding window size of 21 informative sites (half-window size of 10 sites).
`SweepFinder2`: the composite likelihoods were computed on a grid of 800 locations for the selected site (250 nucleotides spacing, four times denser than that used in the original article on `SweepFinder2`).
`VolcanoFinder`: the model 1 was used to compute the composite likelihoods on a grid of 800 locations for the selected site and 13 values for the divergence parameter $D \in \{0.005, 0.01, 0.015, \ldots, 0.065\}$, encompassing the full range of $D$ in the simulations.

*Genome scans for balancing selection.* We compared the performance of `BALLET` and `VolcanoFinder` under three demographic scenarios (Fig. S2.2) inspired by the original `BALLET` article [48]. For each method, the genomic background reference was inferred from a single coalescent simulation of a $2 \times 10^7$ nucleotides under the same demographic scenario.
`BALLET`: the $T2$ test of `BALLET` [48] was used with a sliding window size of 21 informative sites (half-window size of 10 informative sites).
`VolcanoFinder`: the model 1 was used to compute the composite likelihoods on a grid of 500 locations for the selected site and 13 values for the divergence parameter $D \in \{0.001, \ldots, 0.013\}$, enabling to encompass the full range of starting time for the selection $T_s$ in the simulations.

**Power analysis**. For all methods, the maximum LR value in a simulated sequence of 200kb was used as a test statistics.

*Rejection rates.* The rejection rate of a method for a given false positive rate $FPR$ (up to 0.05) was estimated as the proportion of the non-neutral simulations (among 1 000 non-neutral replicates) leading to a test statistics exceeding the $(1 - FPR)$ quantile of the null distribution (estimated from 10 000 neutral replicates). Because introgression sweeps with a low selection coefficient ($2Ns = 100$) mainly altered the site frequency spectrum within 10 kb from the selected site (see Fig. S2.3 and Fig. S2.4), we also computed rejection rates based on the maximum LR value in the central 20 kb in this case.

*Probability of detection of an introgression sweep in a genome scan.* Whole genome scan usually look for outliers among genome-wide data and selection at a locus is detected if the LR value at this locus ranks among the genome-wide highest peaks. We used the following procedure to mimic such a study: the genome-wide null-distributions of the LR values were obtained from the reference 10 000 neutral replicates of 200 kb (leading to 2 Gb genomes). For each neutral replicate 800 LR values were retained (all values for `VolcanoFinder` and `SweepFinder2` and the highest LR value in 800 non-overlapping window of 250 nt for BALLET) leading to $8 \times 10^6$ LR values for the whole genome. The probability to detect an introgression sweep in a genome scan considering a set of top-$X$ candidates was computed as the proportion of the non-neutral replicates (estimated from 1 000 replicates) leading to a maximum LR value (in a single replicate) higher than the $X$th genome-wide neutral highest LR peak.

**Fig. S2.3 Volcano patterns caused by a hard introgression sweep**. Average nucleotide diversity (Tajima's $\hat{\theta}_\pi$, [139]) in non-overlapping windows of 400 nucleotides in the simulated 200 kb alignments involving a hard introgression sweep. The selection strength is $2Ns = 100$ (left) or $2Ns = 1\,000$ (right). The age of the split between the donor and recipient populations is (from top to bottom) $T_d = 1$, 2.5, 4, 5.5 units of $4N$ generations ($D/\theta = 3$, 6, 9, 12). The ending time of the selective sweep is $T_s = 0$ (red), $T_s = 0.1$ (green), $T_s = 0.25$ (blue), $T_s = 0.5$ (black) units of $4N$ generations before sampling. The coloured horizontal lines indicate the background polymorphism level in all cases. For the lowest selection strength only the central 40 kb region is shown.

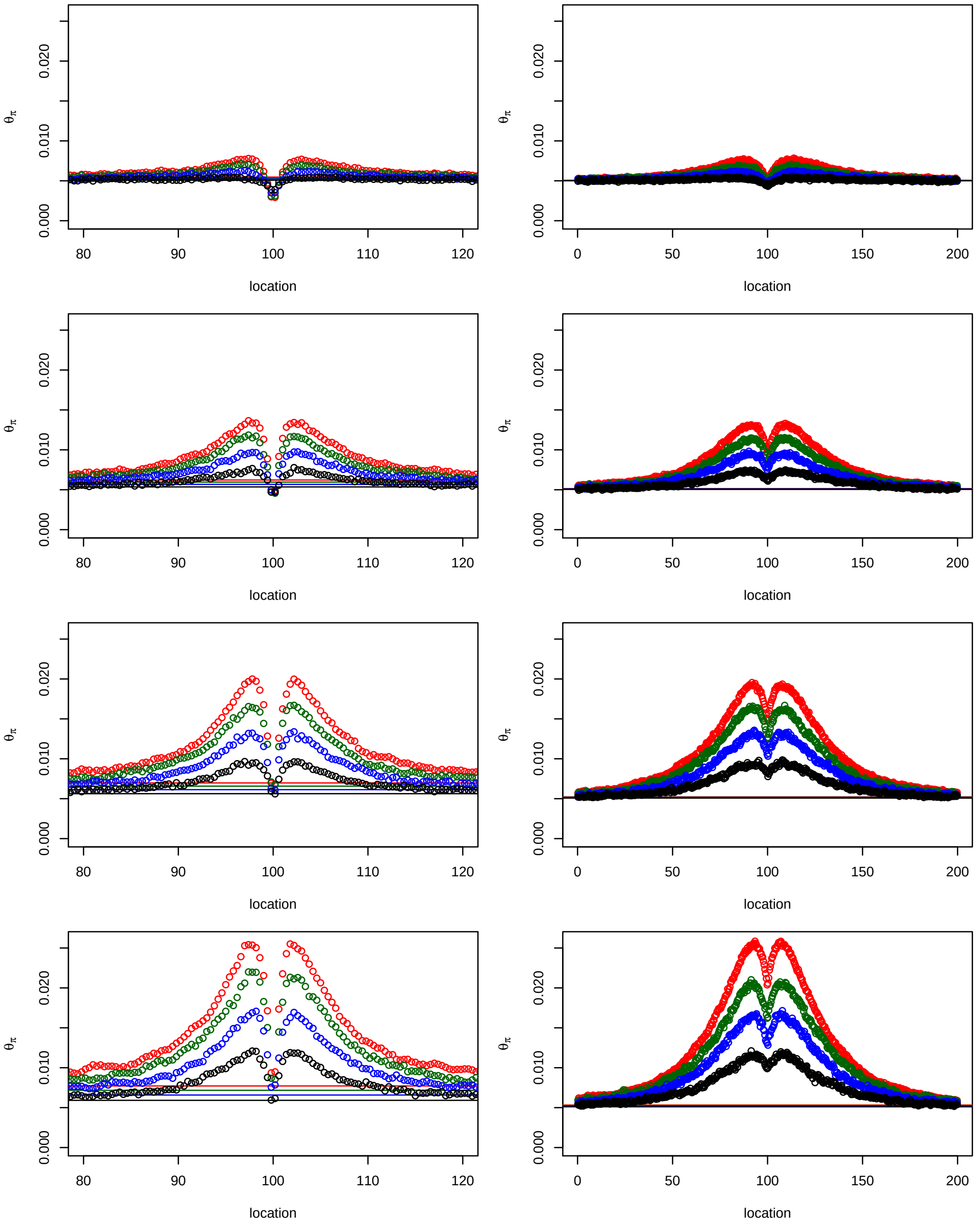

**Fig. S2.4 Volcano patterns caused by a soft introgression sweep**. Average nucleotide diversity (Tajima's $\hat{\theta}_\pi$ [139]) in non-overlapping windows of 400 nucleotides in the simulated 200 kb alignments involving a soft introgression sweep. The selection strength is $2Ns = 100$ (left) or $2Ns = 1\,000$ (right). The age of the split between the donor and recipient populations is (from top to bottom) $T_d = 1$, 2.5, 4, 5.5 units of $4N$ generations ($D/\theta = 3$, 6, 9, 12). The ending time of the selective sweep is $T_s = 0$ (red), $T_s = 0.1$ (green), $T_s = 0.25$ (blue), $T_s = 0.5$ (black) units of $4N$ generations before sampling. The coloured horizontal lines indicate the background polymorphism level in all cases. For the lowest selection strength only the central 40 kb region is shown.

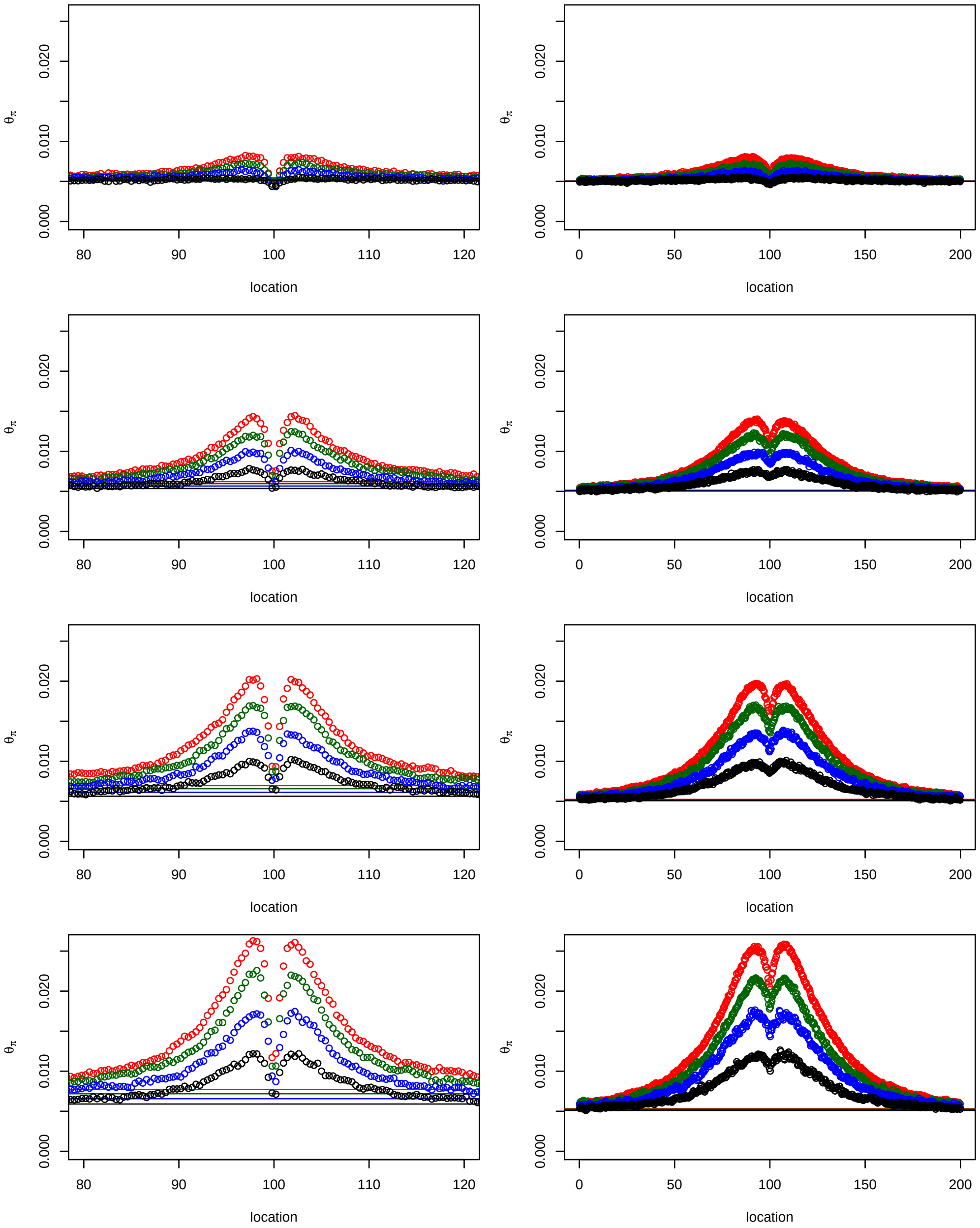

**Fig. S2.5 Performance curves (non admixed background).** Rejection rates of `VolcanoFinder` (blue), `BALLET` (brown) and `SweepFinder2` (green) for an introgression sweep event from a donor species that diverged from the recipient species at (top to bottom) $T_d = 1$, 2.5, 4, 5.5 ($D/\theta = 3$, 6, 9, 12) and a selective sweep that ended (from left to right) $T_s = 0$, 0.1, 0.25, 0.5 units of $4N$ generations in the past. Solid lines: no polymorphism in the donor species (hard introgression sweep). Dashed lines: polymorphism exists in the donor species (possible soft sweep). Dark colour: $2Ns = 1\,000$; light colour: $2Ns = 100$. The upper gray line indicates the expected highest rejection rate given the expected proportion of successful selective sweeps in the sample. Lower gray area: the rejection rate does not exceed the false positive rate. For all three methods, the test statistics is the highest LR value in the simulated 200 000 nucleotides alignment. Analyses involved a neutral non-admixed genomic background as a reference.

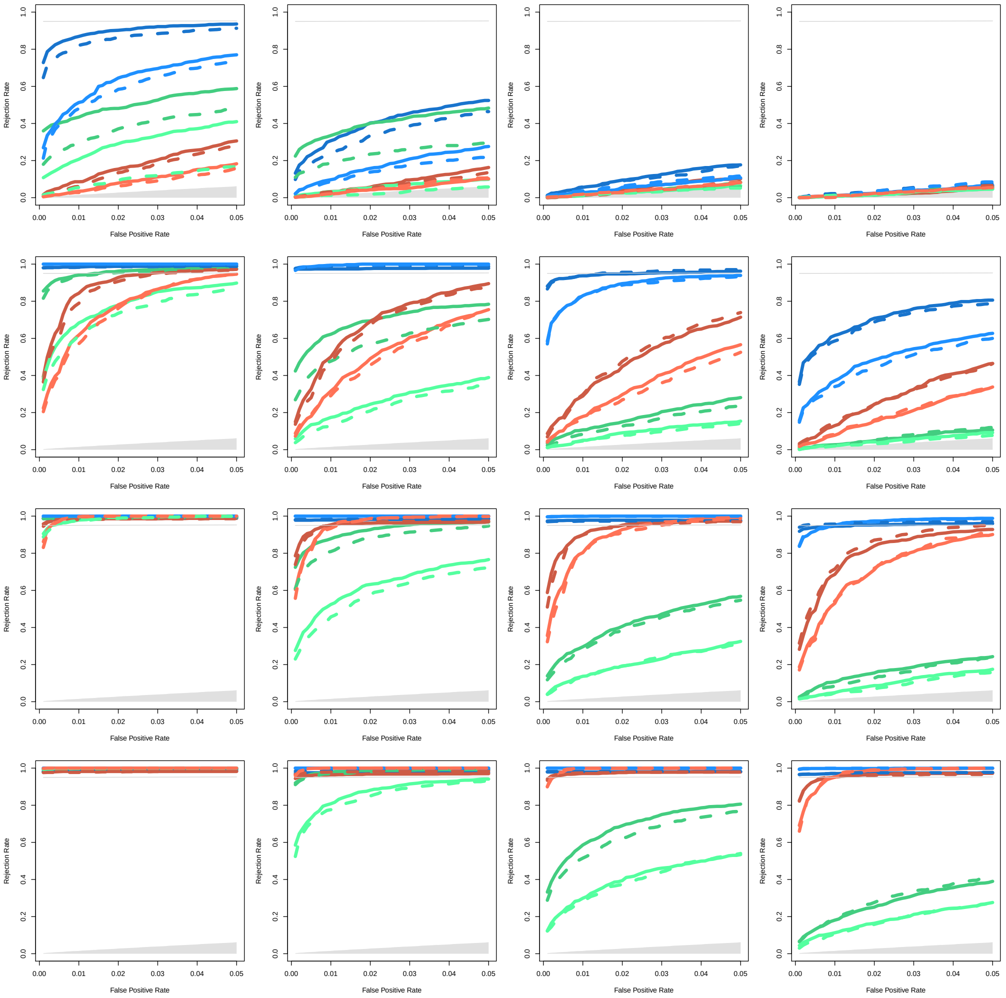

**Fig. S2.6 Performance curves (admixed background).** Rejection rates of `VolcanoFinder` (blue), `BALLET` (brown) and `SweepFinder2` (green) for an introgression sweep event from a donor species that diverged from the recipient species at (top to bottom) $T_d = 1$, 2.5, 4, 5.5 ($D/\theta = 3$, 6, 9, 12) and a selective sweep that ended (from left to right) $T_s = 0$, 0.1, 0.25, 0.5 units of $4N$ generations in the past. Solid lines: no polymorphism in the donor species (hard introgression sweep). Dashed lines: polymorphism exists in the donor species (possible soft sweep). Dark colour: $2Ns = 1\,000$; light colour: $2Ns = 100$. The upper gray line indicates the expected highest rejection rate given the expected proportion of successful selective sweeps in the sample. Lower gray area: the rejection rate does not exceed the false positive rate. For all three methods, the test statistics is the highest LR value in the simulated 200 000 nucleotides alignment. Analyses involved a neutral admixed genomic background with the same level of admixture as a reference.

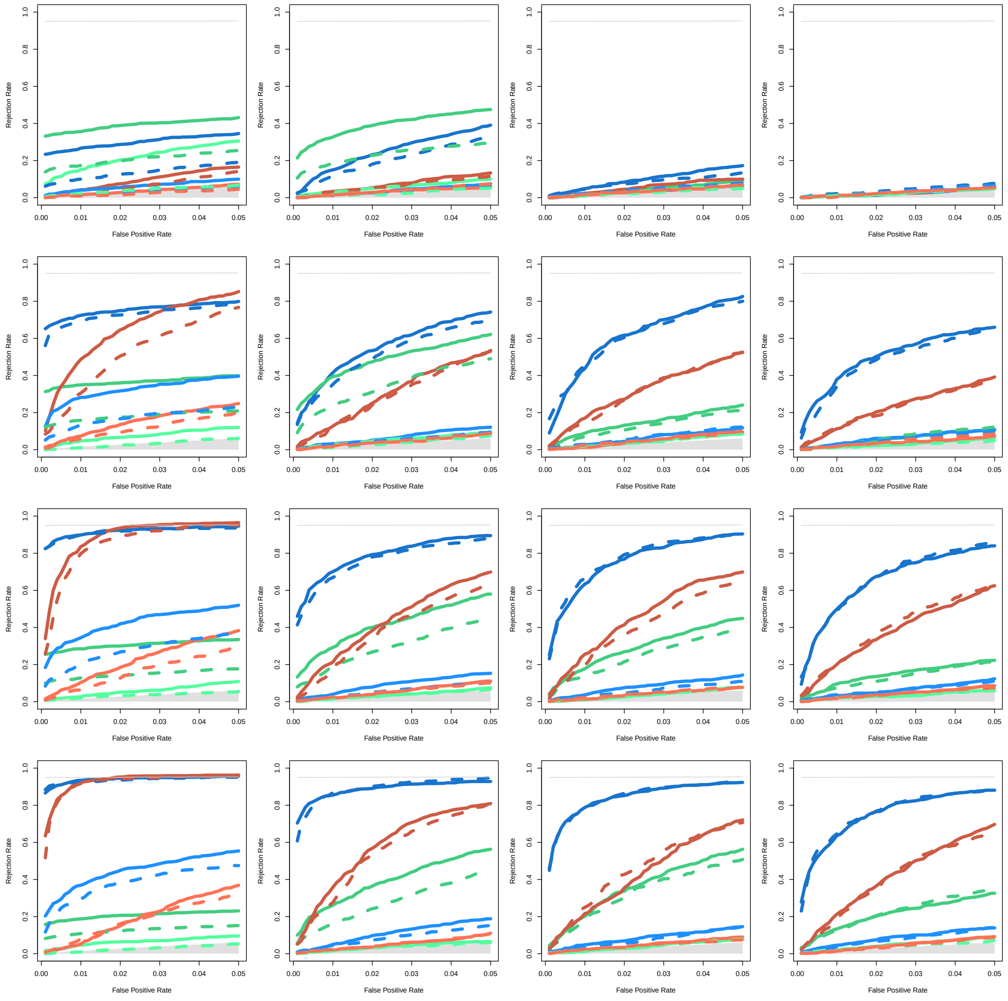

**Fig. S2.7 Performance curves (admixed background and different window sizes).** Rejection rates of `VolcanoFinder` (blue), `BALLET` (brown) and `SweepFinder2` (green) for an introgression sweep event from a donor species that diverged from the recipient species at (top to bottom) $T_d = 1, 2.5, 4, 5.5$ ($D/\theta = 3, 6, 9, 12$) and a selective sweep that ended (from left to right) $T_s = 0, 0.1, 0.25, 0.5$ units of $4N$ generations in the past. Solid lines: no polymorphism in the donor species (hard introgression sweep). Dashed lines: polymorphism exists in the donor species (possible soft sweep). Dark colour: $2Ns = 1\,000$; light colour: $2Ns = 100$. The upper gray line indicates the expected highest rejection rate given the expected proportion of succesfull selective sweeps in the sample. Lower gray area: the rejection rate does not exceed the false positive rate. For all three methods, the test statistics is the highest LR value in the simulated 200 000 nucleotides alignment. The test statistics is the highest LR value in the simulated sequence of 200 kb ($2Ns = 1\,000$) or in the central 20 kb ($2Ns = 100$). Analyses involved a neutral admixed genomic background with the same level of admixture as a reference.

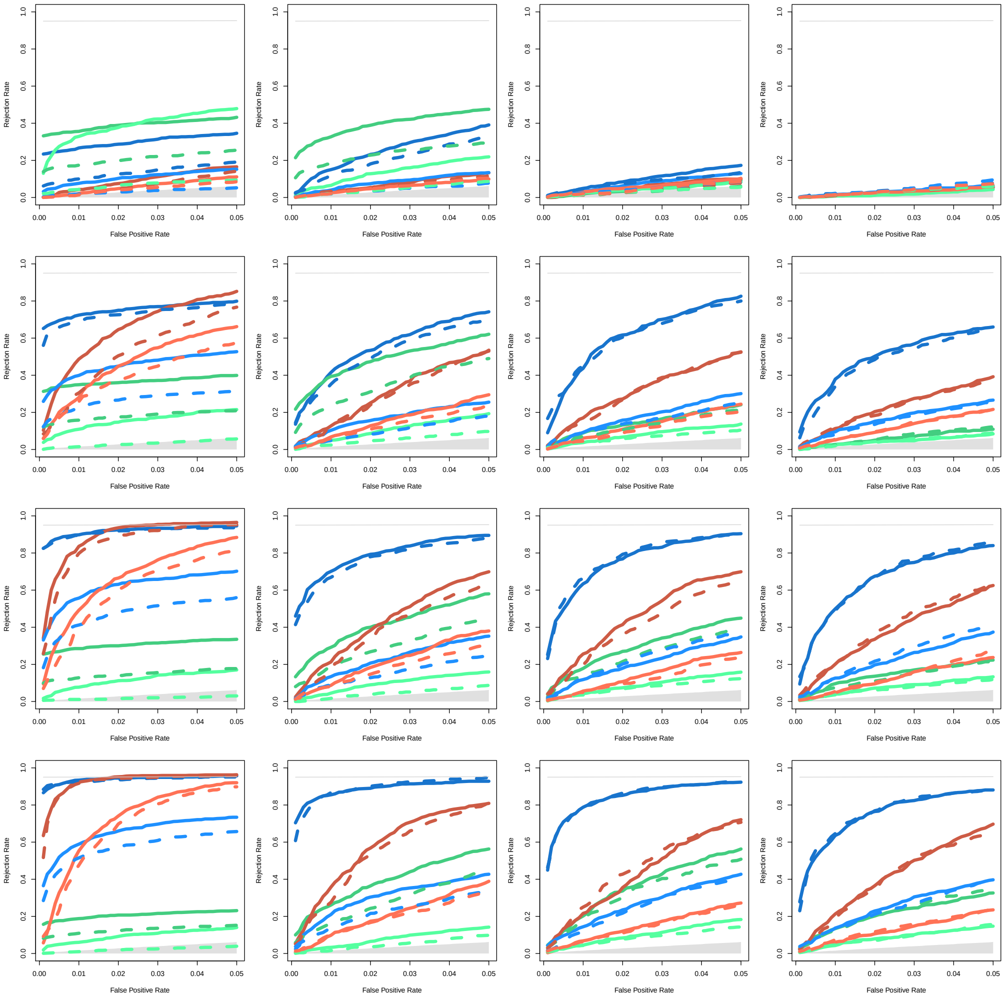

### Text S2.4 Inferred parameters of the selection model.

We assessed the accuracy of VolcanoFinder to infer the position of the selected site, the compound selection parameter $\alpha$, and the divergence with the donor species $D$. For the position of the selected locus, comparisons could be made with the BALLET and SweepFinder methods. We report the observed distributions of the estimated parameters for each parameter set in the case of a hard introgression sweep. In these distributions we highlight values that lead to a significant CLR test (using an admixed background and a 20 kb window size for $2Ns = 100$ and a 200 kb window size for $2Ns = 1\,000$ as in Fig. S2.7).

#### Location of the selected locus as inferred by genome scan methods.

Distributions for the location of the selected site as inferred by the highest CLR value are shown on supp. Fig. S2.8 to Fig. S2.13. VolcanoFinder and SweepFinder2 use information from the valley of reduced heterozygosity in the center of the sweep region and locate the target of selection more accurately than BALLET, which tries to fit a balancing selection model to the data and thus tends to locate the target of selection in the flanking regions where the polymorphism to divergence ratio is higher. For older introgression sweeps ($T_s \geqslant 0.25$) the accuracy of all methods decreases.

#### Parameters of the introgression sweep as inferred by VolcanoFinder.

The distributions of the scaled divergence parameter $\hat{D}/\theta$ inferred from the location with the highest CLR value are shown on Fig. S2.14 to Fig. S2.15. As expected from the analytical analysis, VolcanoFinder tends to underestimate $D$. Unsurprisingly, the mean of the distribution of estimated $\hat{D}$ tends to decrease for older introgression sweeps that typically lead to less pronounced volcano shapes (see Fig. S2.3). The variance of the distribution of $\hat{D}$ also tends to increase with increasing age of the introgression sweep, probably because our model only considers very recent sweeps.

The distributions of the selection strength inferred parameter $-\log_{10}(\hat{\alpha})$ from the location with the highest CLR value are shown on Fig. S2.16 to Fig. S2.17. $-\log_{10}(\alpha)$ seems to be relatively accurately estimated for recent introgression sweeps whereas it might be underestimated in the case of old introgression sweeps that typically lead to narrower volcano shapes (see Fig. S2.3).

**Fig. S2.8 Location of the maximum LR inferred by `VolcanoFinder` ($2Ns = 1\,000$).** Location of the highest LR inferred by `VolcanoFinder` for a hard introgression sweep event with selection coefficient $2Ns = 1\,000$. The donor species diverged from the recipient species at (from top to bottom) $T_d = 1$, 2.5, 4, 5.5 ($D/\theta = 3$, 6, 9, 12) and the selective sweep ended (from left to right) $T_s = 0$, 0.1, 0.25, 0.5 units of $4N$ generations in the past. The coloured and gray parts indicate significant and non-significant test as shown in Fig. S2.7.

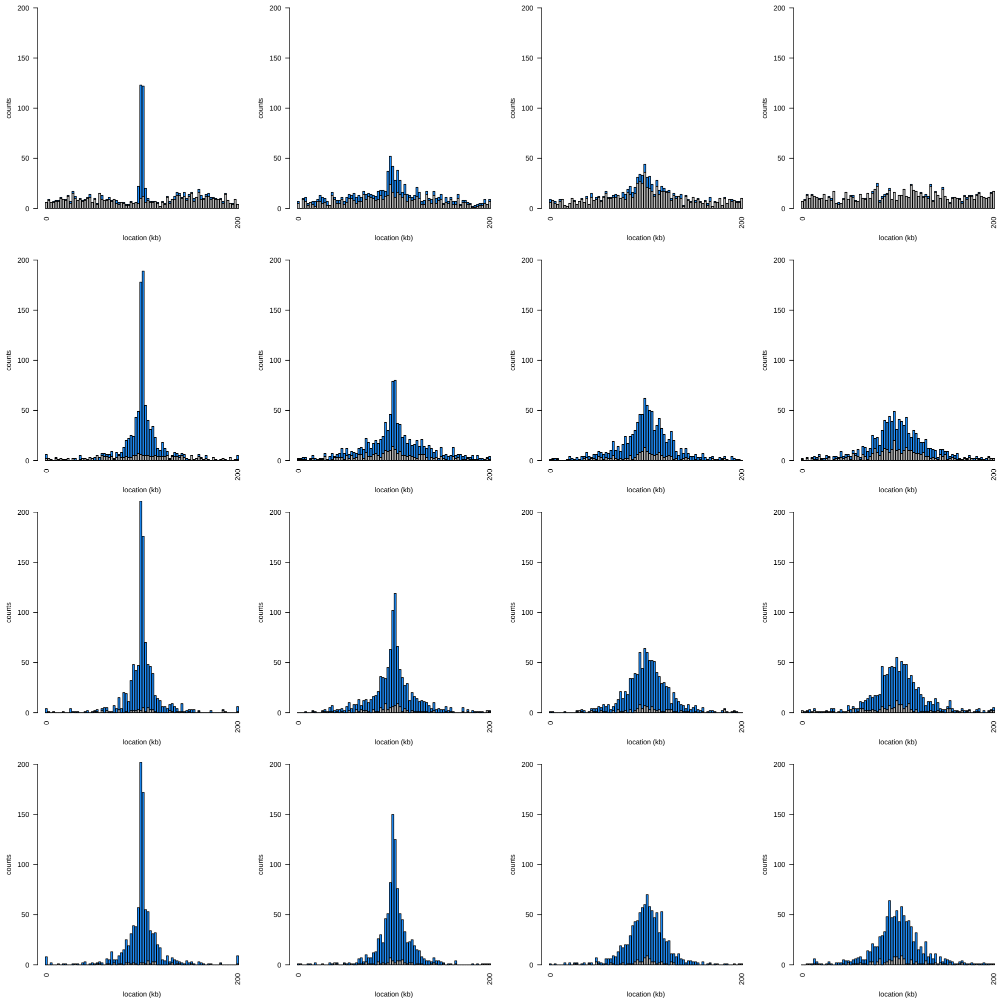

**Fig. S2.9 Location of the maximum LR inferred by `VolcanoFinder` ($2Ns = 100$).** Location of the highest LR inferred by `VolcanoFinder` for a hard introgression sweep event with selection coefficient $2Ns = 100$. The donor species diverged from the recipient species at (from top to bottom) $T_d = 1$, 2.5, 4, 5.5 ($D/\theta = 3$, 6, 9, 12) and the selective sweep ended (from left to right) $T_s = 0$, 0.1, 0.25, 0.5 units of $4N$ generations in the past. The coloured and gray parts indicate significant and non-significant test as shown in Fig. S2.7. Only the central part of the simulated region is shown.

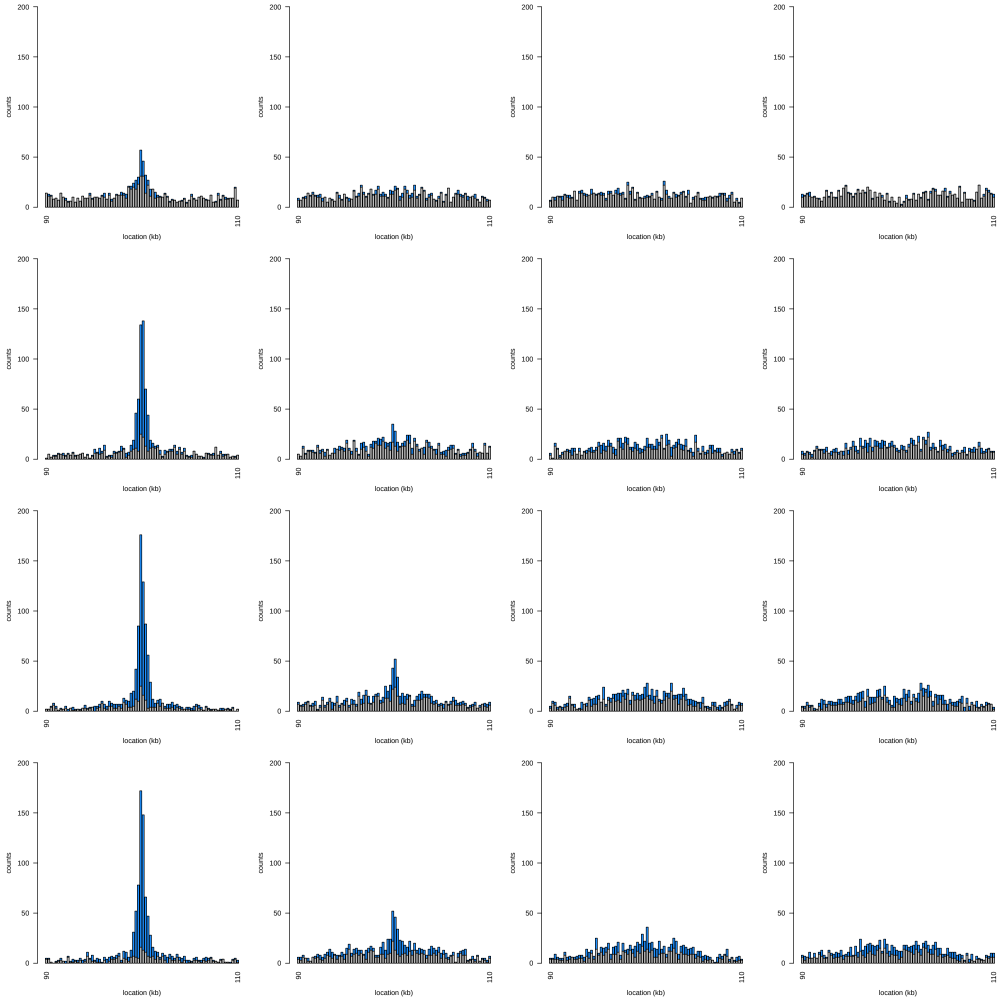

**Fig. S2.10 Location of the maximum LR inferred by `SweepFinder2` ($2Ns = 1\,000$).** Location of the highest LR inferred by `SweepFinder2` for a hard introgression sweep event with selection coefficient $2Ns = 1\,000$. The donor species diverged from the recipient species at (from top to bottom) $T_d = 1$, 2.5, 4, 5.5 ($D/\theta = 3$, 6, 9, 12) and the selective sweep ended (from left to right) $T_s = 0$, 0.1, 0.25, 0.5 units of $4N$ generations in the past. The coloured and gray parts indicate significant and non-significant test as shown in Fig. S2.7.

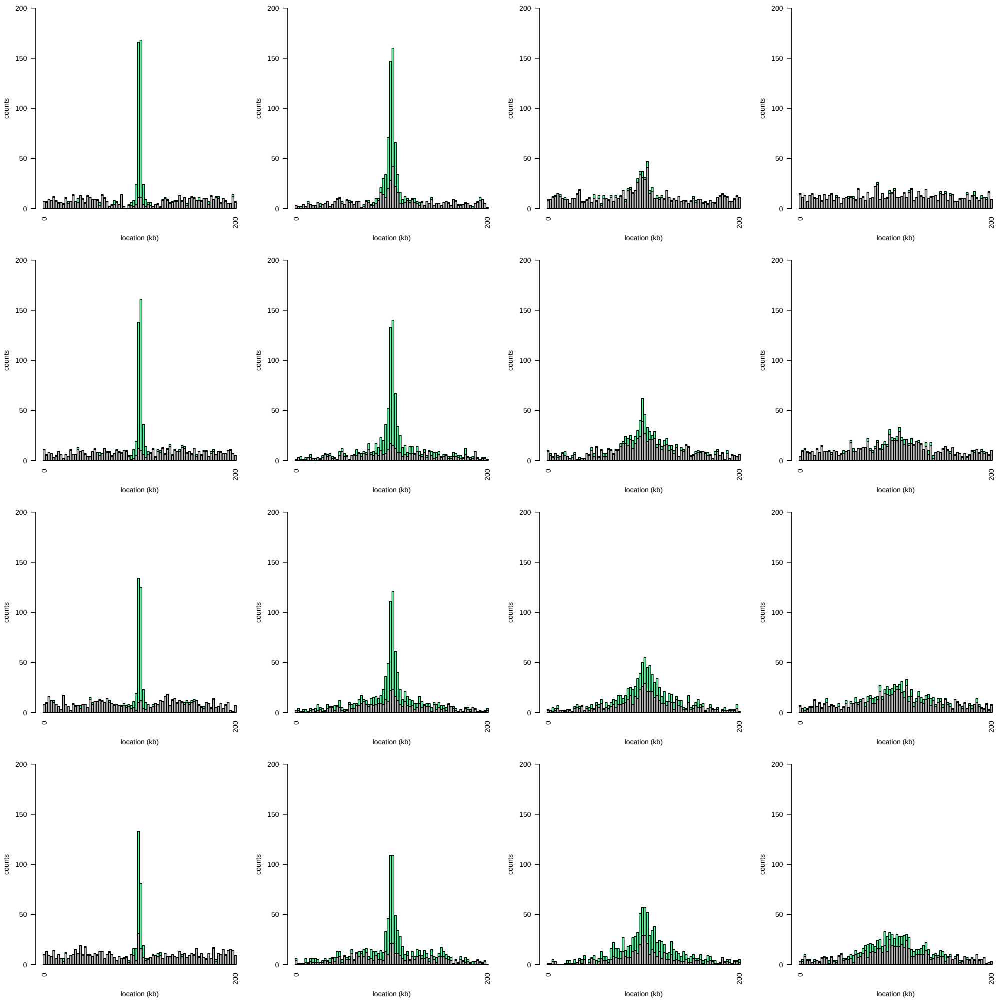

**Fig. S2.11 Location of the maximum LR inferred by `SweepFinder2` ($2Ns = 100$).** Location of the highest LR inferred by `SweepFinder2` for a hard introgression sweep event with selection coefficient $2Ns = 100$. The donor species diverged from the recipient species at (from top to bottom) $T_d = 1$, 2.5, 4, 5.5 ($D/\theta = 3$, 6, 9, 12) and the selective sweep ended (from left to right) $T_s = 0$, 0.1, 0.25, 0.5 units of $4N$ generations in the past. The coloured and gray parts indicate significant and non-significant test as shown in Fig. S2.7. Only the central part of the simulated region is shown.

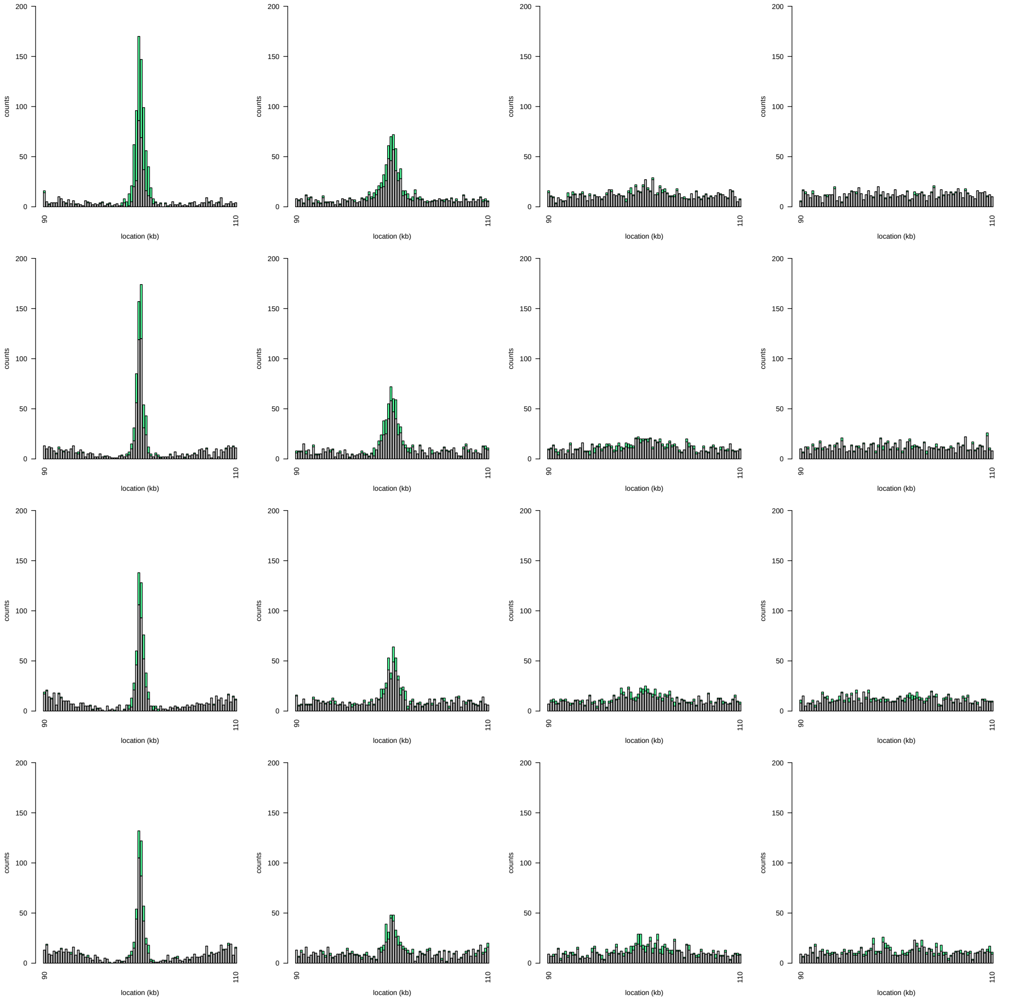

**Fig. S2.12 Location of the maximum LR inferred by BALLET ($2Ns = 1\,000$).** Location of the highest LR inferred by BALLET for a hard introgression sweep event with selection coefficient $2Ns = 1\,000$. The donor species diverged from the recipient species at (from top to bottom) $T_d = 1$, 2.5, 4, 5.5 ($D/\theta = 3$, 6, 9, 12) and the selective sweep ended (from left to right) $T_s = 0$, 0.1, 0.25, 0.5 units of $4N$ generations in the past. The coloured and gray parts indicate significant and non-significant test as shown in Fig. S2.7.

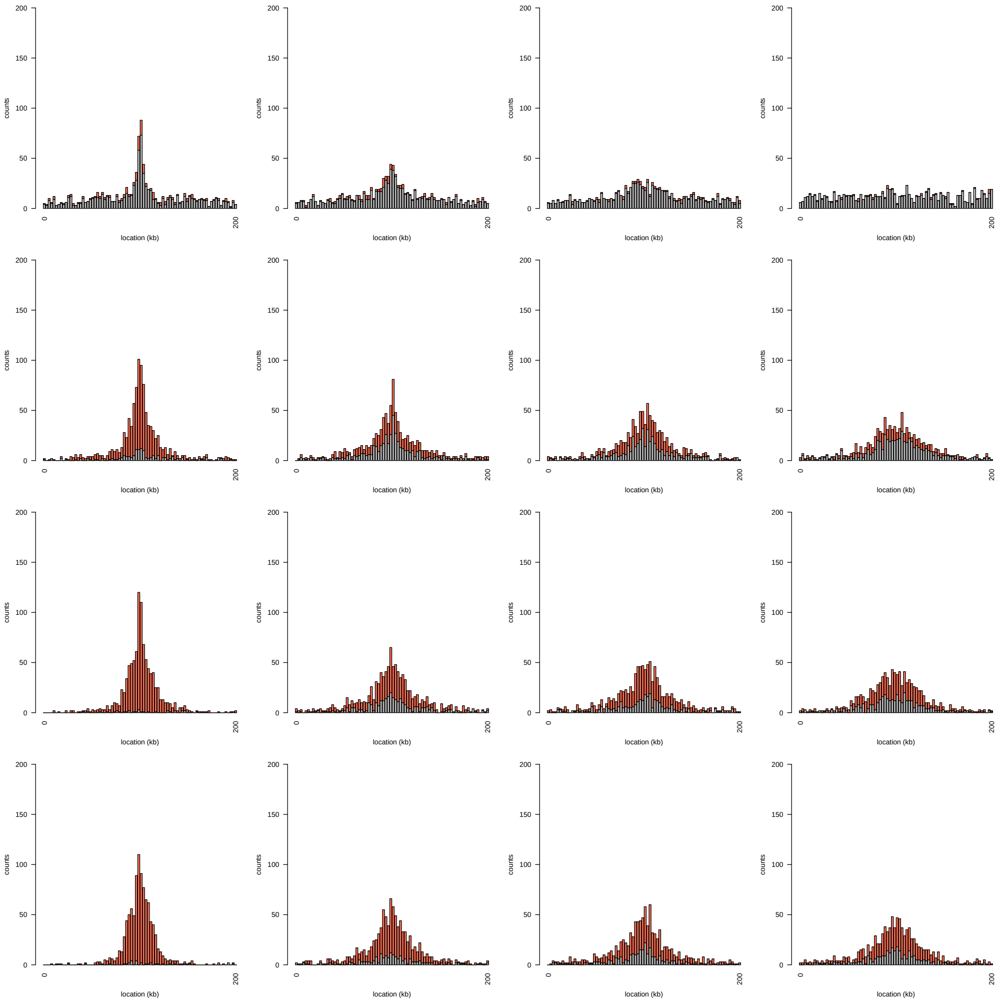

**Fig. S2.13 Location of the maximum LR inferred by BALLET ($2Ns = 100$).** Location of the highest LR inferred by BALLET for a hard introgression sweep event with selection coefficient $2Ns = 100$. The donor species diverged from the recipient species at (from top to bottom) $T_d = 1$, 2.5, 4, 5.5 ($D/\theta = 3$, 6, 9, 12) and the selective sweep ended (from left to right) $T_s = 0$, 0.1, 0.25, 0.5 units of $4N$ generations in the past. The coloured and gray parts indicate significant and non-significant test as shown in Fig. S2.7. Only the central part of the simulated region is shown.

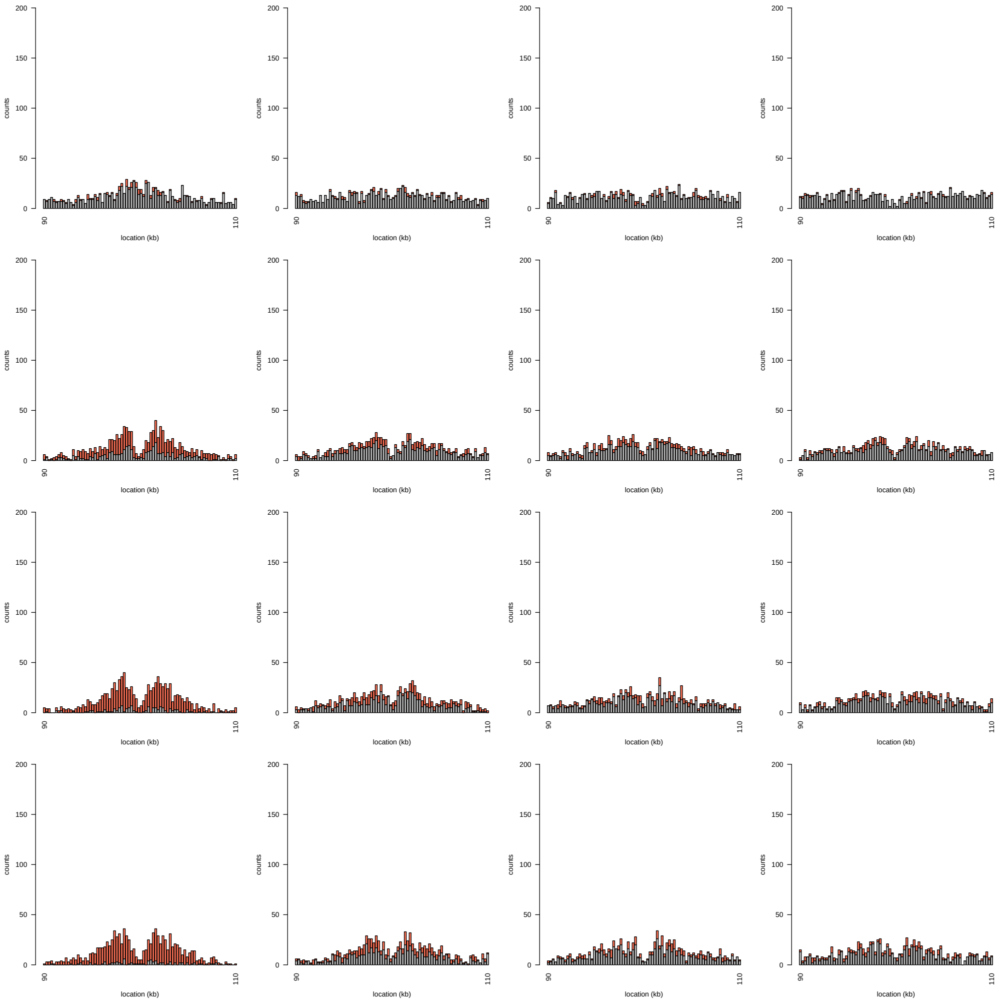

**Fig. S2.14 Divergence from the donor species inferred by `VolcanoFinder` ($2Ns = 1\,000$, hard introgression sweeps)**. Estimated scaled divergence parameter $\frac{\hat{D}}{\theta}$ at the location with the highest LR inferred by `VolcanoFinder` for a hard introgression sweep event with selection coefficient $2Ns = 1\,000$. The donor species diverged from the recipient species at (from top to bottom) $T_d = 1$, 2.5, 4, 5.5 ($D/\theta = 3$, 6, 9, 12) and the selective sweep ended (from left to right) $T_s = 0$, 0.1, 0.25, 0.5 units of $4N$ generations in the past. The coloured and gray parts indicate significant and non-significant test as shown in Fig. S2.7. A vertical red line indicates the true value used in the simulations.

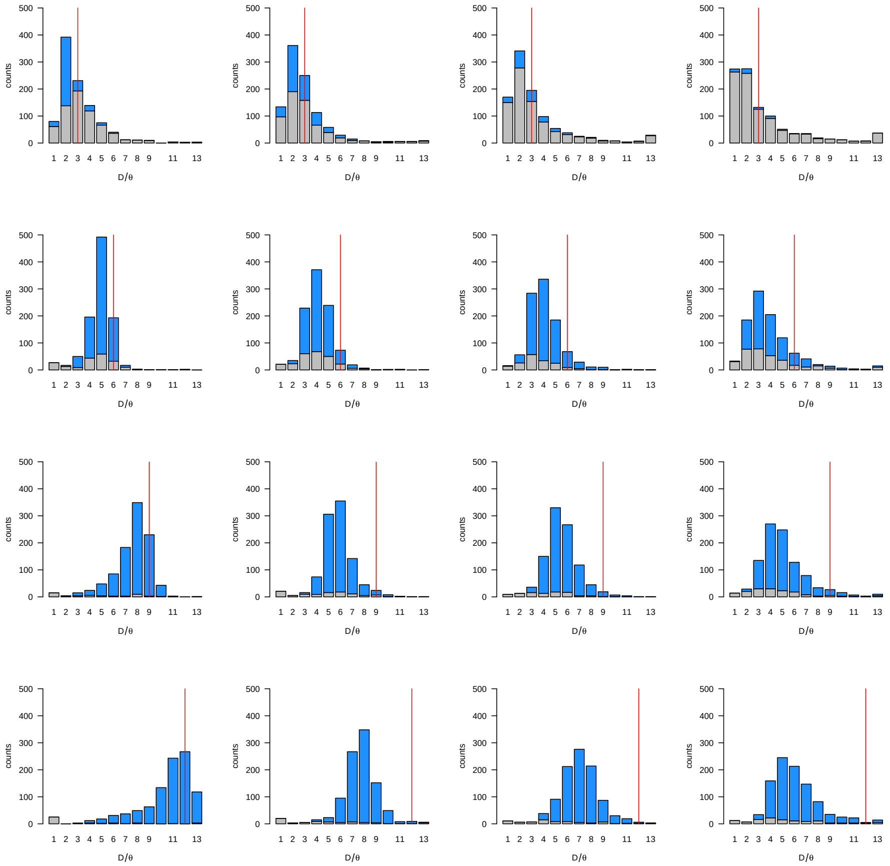

**Fig. S2.15 Divergence from the donor species inferred by `VolcanoFinder` ($2Ns = 100$, hard introgression sweeps).** Estimated scaled divergence parameter $\frac{\hat{D}}{\theta}$ for the location with the highest LR inferred by `VolcanoFinder` for a hard introgression sweep event with selection coefficient $2Ns = 100$. The donor species diverged from the recipient species at (from top to bottom) $T_d = 1$, 2.5, 4, 5.5 ($D/\theta = 3$, 6, 9, 12) and the selective sweep ended (from left to right) $T_s = 0$, 0.1, 0.25, 0.5 units of $4N$ generations in the past. The coloured and gray parts indicate significant and non-significant test as shown in Fig. S2.7. A vertical red line indicates the true value used in the simulations.

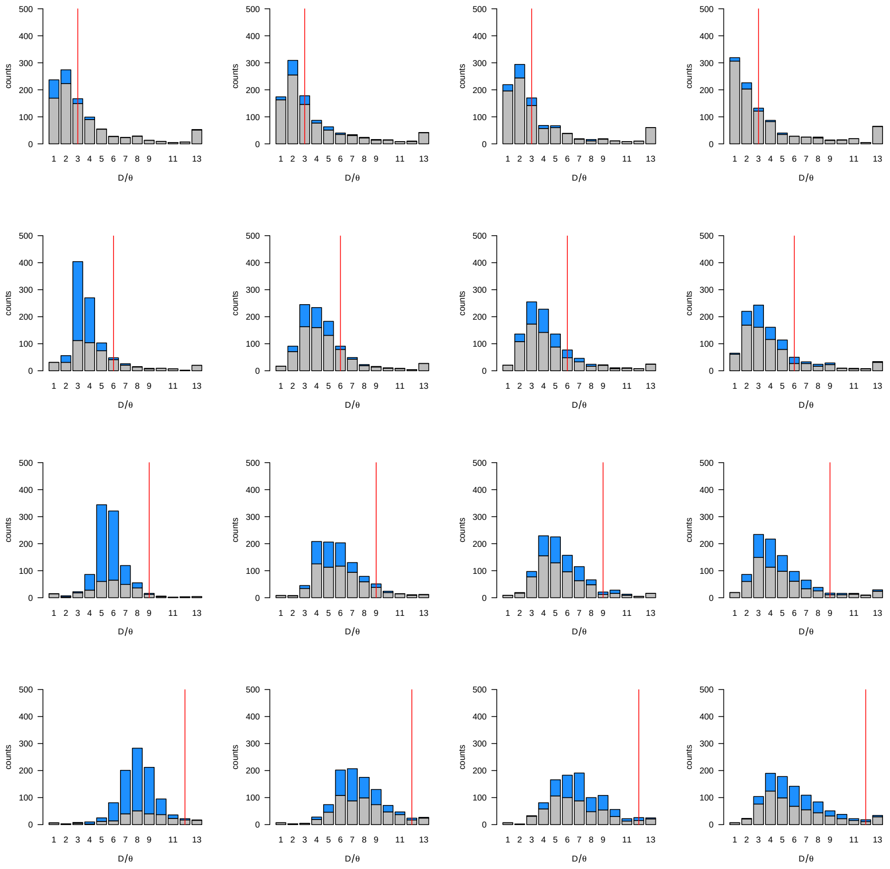

**Fig. S2.16 Selection strength inferred by `VolcanoFinder` ($2Ns = 1\,000$, hard introgression sweeps)**. Estimated scaled selection parameter $-\log_{10}(\hat{\alpha})$ at the location with the highest LR inferred by `VolcanoFinder` for a hard introgression sweep event with selection coefficient $2Ns = 1\,000$. The donor species diverged from the recipient species at (from top to bottom) $T_d = 1$, 2.5, 4, 5.5 ($D/\theta = 3$, 6, 9, 12) and the selective sweep ended (from left to right) $T_s = 0$, 0.1, 0.25, 0.5 units of $4N$ generations in the past. The coloured and gray parts indicate significant and non-significant test as shown in Fig. S2.7. A vertical red line indicates the true value used in the simulations.

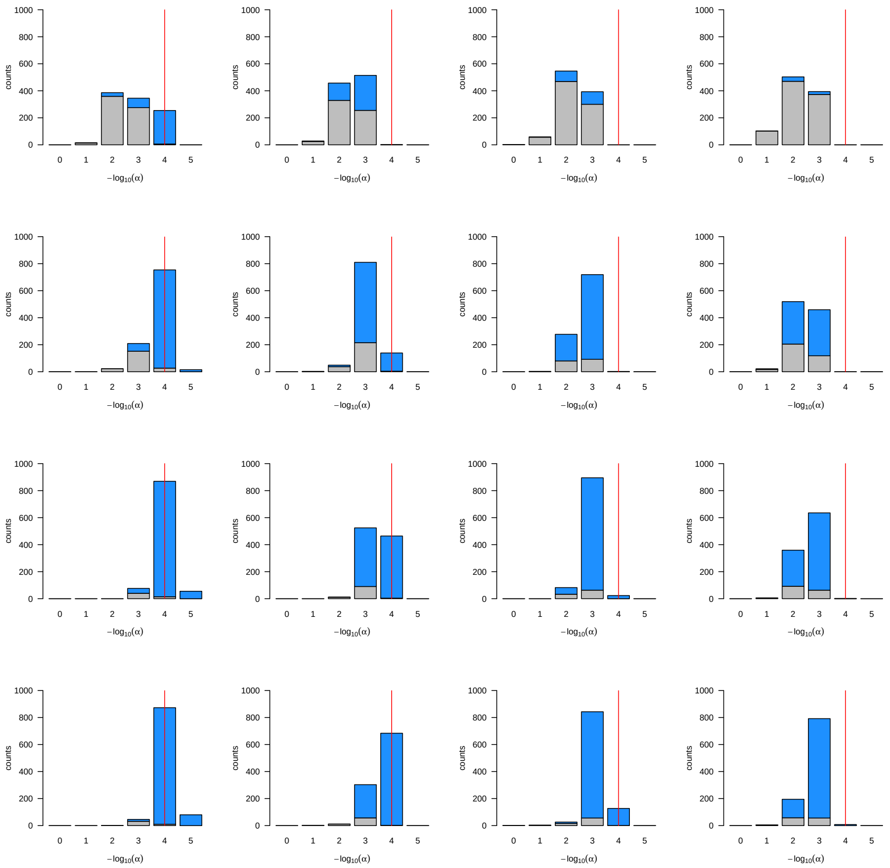

**Fig. S2.17 Selection strength inferred by `VolcanoFinder` ($2Ns = 100$, hard introgression sweeps).** Estimated scaled selection parameter $-\log_{10}(\hat{\alpha})$ for the location with the highest LR inferred by `VolcanoFinder` for a hard introgression sweep event with selection coefficient $2Ns = 100$. The donor species diverged from the recipient species at (from top to bottom) $T_d = 1$, 2.5, 4, 5.5 ($D/\theta = 3$, 6, 9, 12) and the selective sweep ended (from left to right) $T_s = 0$, 0.1, 0.25, 0.5 units of $4N$ generations in the past. The coloured and gray parts indicate significant and non-significant test as shown in Fig. S2.7. A vertical red line indicates the true value used in the simulations.

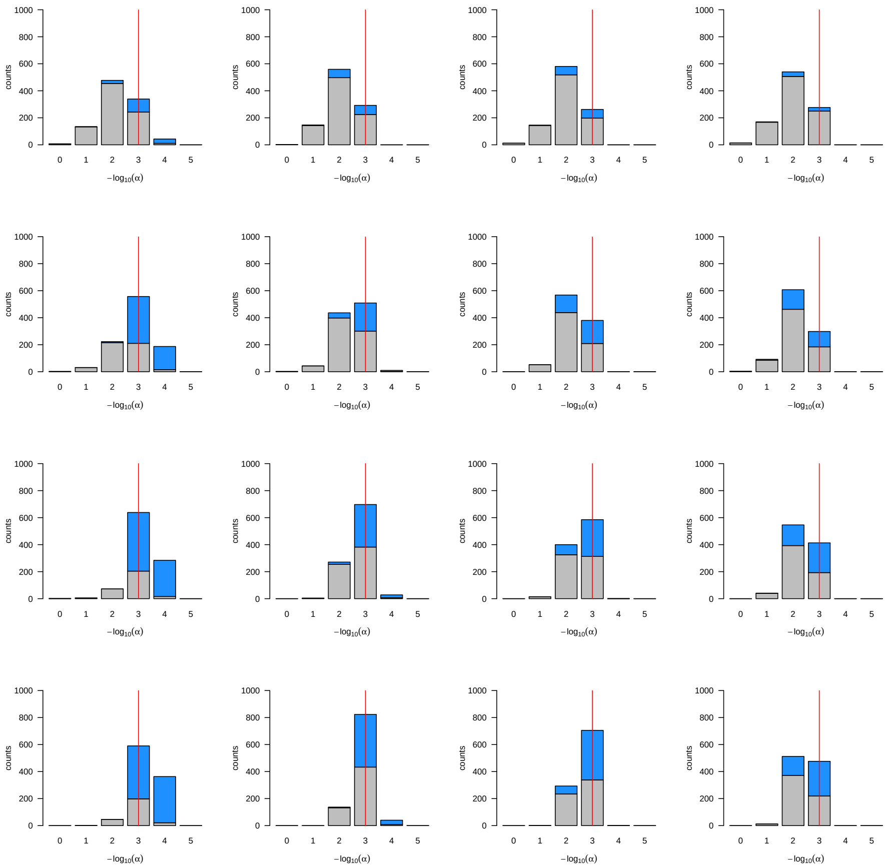

**Fig. S2.18 Performance curves to detect balancing selection**. Rejection rates of `BALLET` (solid lines) and `VolcanoFinder` (dashed lines) for different starting time of balancing selection $T_s$ under three demographic scenarios (see Fig. S2.2). Black: constant population size; blue: population growth; red: bottleneck. Gray area: the rejection rate is not higher than the false positive rate. In all cases, the test statistics is the highest LR value in the simulated 200 kb alignment.

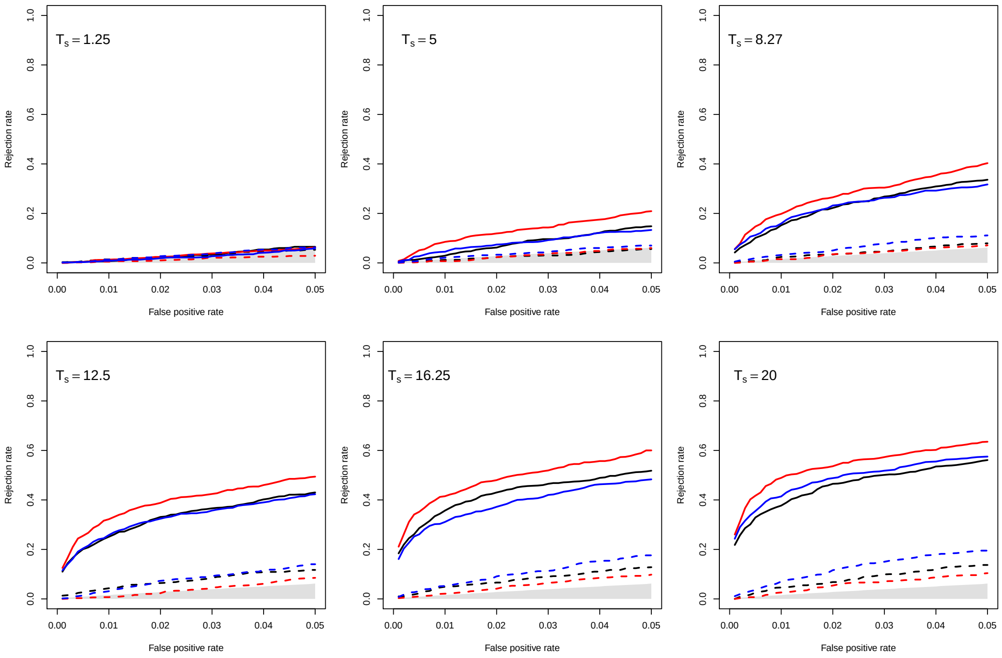

**Table S3.1 Candidate peaks ranked by the maximum log likelihood ratios in the `VolcanoFinder` scan of the European (CEU) sample.**

| Chr. | Peak Position | L.R. | $-\log_{10}\hat{\alpha}$ | $\hat{D}$ | Nearest Gene(s) | RefSeq ID |
|---|---|---|---|---|---|---|
| 22 | 36 556 023 | 102.4 | 3.42 | 0.0038 | *APOL3*, *APOL4* | NM_145640.2, NM_030643.4 |
| 8 | 42 558 168 | 80.4 | 3.67 | 0.0023 | *CHRNB3*, *CHRNA6* | NM_001347717.1, NM_004198.3 |
| 2 | 223 953 190 | 55.5 | 3.21 | 0.0061 | *KCNE4* | NM_080671.3 |
| 14 | 81 512 174 | 51.5 | 3.73 | 0.0023 | *TSHR* | NM_001018036.2 |
| 21 | 19 243 059 | 50.5 | 3.68 | 0.0015 | *CHODL*, *CHODL-AS1* | NM_001204177.1, NR_024354.1 |
| 2 | 24 626 190 | 36.9 | 3.88 | 0.0015 | *ITSN2* | NM_001348181.1 |
| 2 | 223 924 190 | 36.7 | 3.58 | 0.0030 | *KCNE4* | NM_080671.3 |
| 3 | 182 989 072 | 35.4 | 3.48 | 0.0023 | *MCF2L2*, *B3GNT5* | NM_015078.3, NM_032047.4 |
| 2 | 122 044 190 | 35.2 | 3.50 | 0.0015 | *TFCP2L1* | NM_014553.2 |
| 2 | 172 059 190 | 33.7 | 3.49 | 0.0023 | *TLK1* | NM_012290.4 |
| 7 | 73 935 618 | 32.8 | 3.31 | 0.0030 | *GTF2IRD1* | NM_005685.3 |
| 2 | 12 035 190 | 29.6 | 3.35 | 0.0023 | – | – |
| 16 | 83 231 010 | 29.5 | 3.19 | 0.0015 | *CDH13* | NM_001220491.1 |
| 3 | 127 624 072 | 29.2 | 3.49 | 0.0023 | *KBTBD12* | NM_207335.2 |
| 11 | 44 436 084 | 29.0 | 3.30 | 0.0015 | – | – |
| 12 | 71 969 102 | 28.5 | 3.12 | 0.0038 | *LGR5*, *ZFC3H1* | NM_001277227.1, NM_144982.4 |
| 19 | 17 289 015 | 27.9 | 3.20 | 0.0023 | *MYO9B*, *USE1*, *OCEL1* | NM_004145.3, NM_018467.3, NM_024578.2 |
| 20 | 7 596 076 | 27.8 | 3.32 | 0.0023 | – | – |
| 10 | 28 705 072 | 27.4 | 3.47 | 0.0015 | – | – |
| 1 | 232 398 053 | 26.5 | 3.50 | 0.0015 | – | – |
| 3 | 129 080 072 | 25.6 | 3.22 | 0.0023 | *RPL32P3*, *H1FX*, *H1FX-AS1*, *SNORA7B*, *EFCAB12* | NR_003111.2, NM_006026.3, NR_026991.1, NR_002992.2, NM_207307.2 |
| 5 | 154 678 042 | 24.6 | 3.50 | 0.0015 | – | – |
| 13 | 105 399 042 | 24.4 | 3.28 | 0.0030 | – | – |
| 9 | 115 694 060 | 24.4 | 3.29 | 0.0015 | *SLC46A2* | NM_033051.3 |
| 6 | 29 035 112 | 23.9 | 3.12 | 0.0023 | *LOC100129636*, *OR2W1*, *OR2B3*, *OR2J3* | NR_125387.1, NM_030903.3, NM_001005226.2, NM_001005216.3 |
| 9 | 79 675 060 | 23.9 | 3.27 | 0.0015 | *FOXB2* | NM_001013735.1 |
| 6 | 34 052 112 | 23.2 | 3.29 | 0.0023 | *GRM4* | NM_000841.3 |

**Table S3.2 Candidate peaks ranked by the maximum log likelihood ratios in the `VolcanoFinder` scan of the Yoruban (YRI) sample.**

| Chr. | Peak Position | LR | $-\log_{10}\hat{\alpha}$ | $\hat{D}$ | Nearest Gene(s) | Respective RefSeq ID |
|---|---|---|---|---|---|---|
| 19 | 41 473 015 | 45.2 | 3.49 | 0.0020 | *CYP2B7P*, *CYP2B6* | NR_001278.1, NM_000767.4 |
| 1 | 152 102 007 | 37.3 | 3.48 | 0.0030 | *LOC100131107*, *TCHHL1*, *TCHH*, *RPTN* | NM_001310142.1, NM_001008536.1, NM_007113.3, NM_001122965.1 |
| 12 | 59 033 128 | 32.1 | 3.56 | 0.0020 | *LOC101927653*, *LOC100506869* | NR_120452.1, NR_126341.1 |
| 3 | 33 007 016 | 32.0 | 3.48 | 0.0030 | *CCR4*, *GLB1* | NM_005508.4, NM_001079811.2 |
| 2 | 170 442 117 | 28.7 | 3.25 | 0.0020 | *FASTKD1*, *PPIG* | NM_001322046.1, NM_004792.2 |
| 2 | 235 174 117 | 25.1 | 3.09 | 0.0020 | – | – |
| 4 | 101 771 036 | 22.9 | 3.53 | 0.0020 | – | – |
| 4 | 78 105 036 | 22.2 | 3.31 | 0.0030 | *CCNG2* | NM_004354.2 |

**Fig. S3.1 Whole-genome Manhattan plot of the maximum likelihood ratio test statistic for the European (CEU) population computed from Model 1 of `VolcanoFinder` on data on within-CEU polymorphism and substitutions with respect to chimpanzee, and annotated with the top 18 gene candidates.**

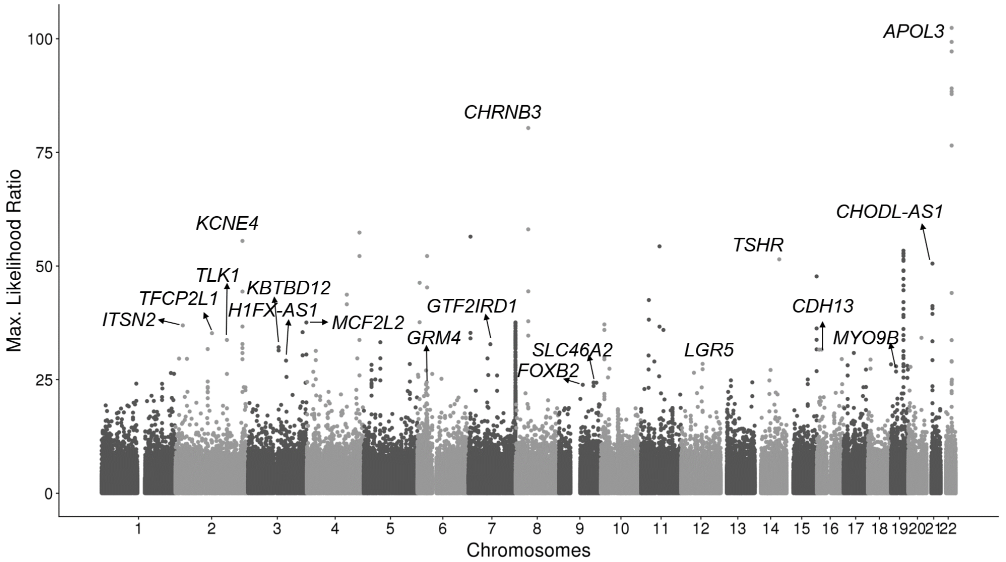

**Fig. S3.2 Whole-genome Manhattan plot of the maximum likelihood ratio test statistic for the Yoruban (YRI) population computed from Model 1 of `VolcanoFinder` on data on within-YRI polymorphism and substitutions with respect to chimpanzee, and annotated with the top 7 gene candidates.**

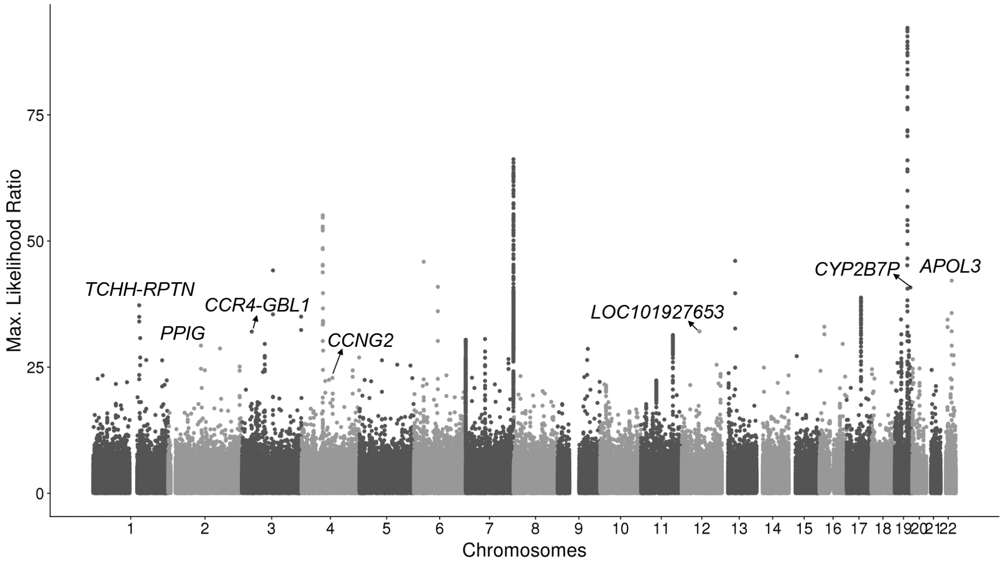

**Fig. S3.3 Introgression sweep signals, parameter estimates, and sequencing properties across the 100 kb region on chromosome 22 covering *APOL* gene cluster in YRI.**

**A.** Likelihood ratio test statistic computed from Model 1 of `VolcanoFinder` on data on within-YRI polymorphism and substitutions with respect to chimpanzee. Horizontal dark gray, medium gray, and light gray bars correspond to regions that were filtered based on Hardy-Weinberg equilibrium (HWE) test. Gene tracts and labels for key genes are depicted below the plot, with the wider bars representing exons. **B.** Values for $\alpha$ and divergence $D$ corresponding to the maximum likelihood estimate of the data. Black line corresponds to $-\ln(\alpha)$ and vertical gray bars correspond to estimated $D$. **C.** Likelihood ratio test statistic computed from $T_2$ of `BALLET` on data on within-YRI polymorphism and substitutions with respect to chimpanzee using windows of 100 (black) or 22 (gray) informative sites on either side of the test site. **D.** Mean pairwise sequence difference ($\hat{\theta}_\pi$) computed in five kb windows centered on each polymorphic site. **E.** Mappability uniqueness scores for 35 nucleotide sequences across the region. **F.** Mean sequencing depth across the 108 YRI individuals as a function of genomic position, with the gray ribbon indicating standard deviation. The background heatmap displays the number of individuals devoid of sequencing reads as a function of genomic position, with darker shades of red indicating a greater number of individuals with no sequencing reads.

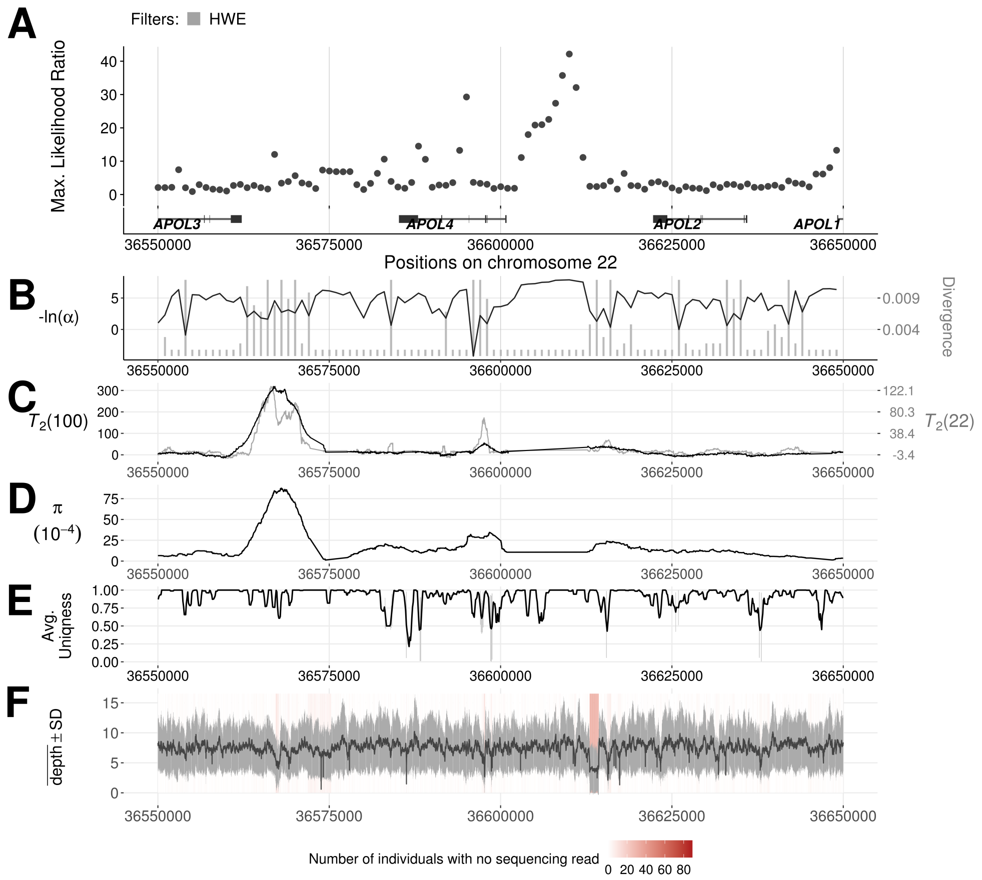

**Fig. S3.4 Introgression sweep signals, parameter estimates, and sequencing properties across the 100 kb region on chromosome 22 covering *APOL4* gene in CEU, matching the same region in YRI.**

**A.** Likelihood ratio test statistic computed from Model 1 of `VolcanoFinder` on data on within-CEU polymorphism and substitutions with respect to chimpanzee. Horizontal dark gray, medium gray, and light gray bars correspond to regions that were filtered based on Hardy-Weinberg equilibrium (HWE) test. Gene tracts and labels for key genes are depicted below the plot, with the wider bars representing exons. **B.** Values for $\alpha$ and divergence $D$ corresponding to the maximum likelihood estimate of the data. Black line corresponds to $-\ln(\alpha)$ and vertical gray bars correspond to estimated $D$. **C.** Likelihood ratio test statistic computed from $T_2$ of `BALLET` on data on within-CEU polymorphism and substitutions with respect to chimpanzee using windows of 100 (black) or 22 (gray) informative sites on either side of the test site. **D.** Mean pairwise sequence difference ($\hat{\theta}_\pi$) computed in five kb windows centered on each polymorphic site. **E.** Mappability uniqueness scores for 35 nucleotide sequences across the region. **F.** Mean sequencing depth across the 108 YRI individuals as a function of genomic position, with the gray ribbon indicating standard deviation. The background heatmap displays the number of individuals devoid of sequencing reads as a function of genomic position, with darker shades of red indicating a greater number of individuals with no sequencing reads.

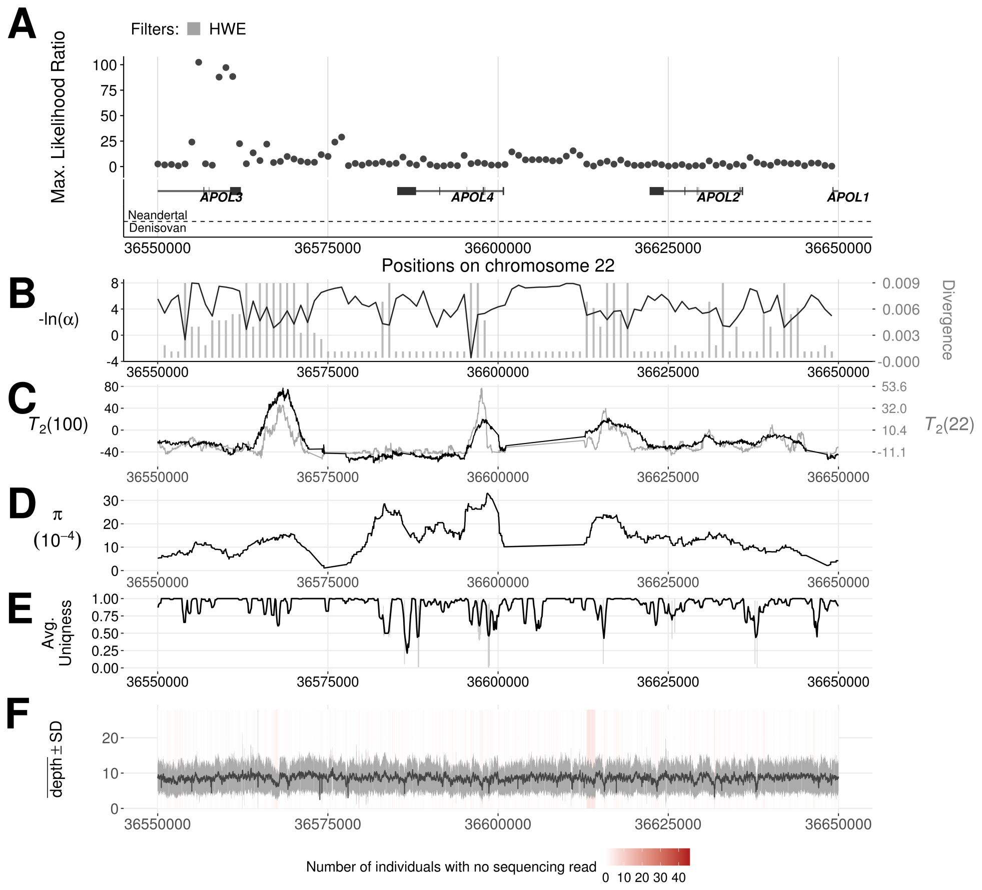

**Fig. S3.5 Introgression sweep signals, parameter estimates, and sequencing properties across the one Mb region on chromosome 7 covering the *PTPRN2* gene region in YRI.**

**A.** Likelihood ratio test statistic computed from Model 1 of `VolcanoFinder` on data on within-YRI polymorphism and substitutions with respect to chimpanzee. Horizontal dark gray and light gray bars correspond to regions that were filtered based on either mean CRG score or mean CRG score and proximity to a telomere, respectively. Gene tracts and labels for key genes are depicted below the plot, with the wider bars representing exons. **B.** Values for $\alpha$ and divergence $D$ corresponding to the maximum likelihood estimate of the data. Black line corresponds to $-\ln(\alpha)$ and vertical gray bars correspond to estimated $D$. **C.** Likelihood ratio test statistic computed from $T_2$ of `BALLET` on data on within-YRI polymorphism and substitutions with respect to chimpanzee using windows of 100 (black) or 22 (gray) informative sites on either side of the test site. **D.** Mean pairwise sequence difference ($\hat{\theta}_\pi$) computed in five kb windows centered on each polymorphic site. **E.** Mappability uniqueness scores for 35 nucleotide sequences across the region. **F.** Mean sequencing depth across the 108 YRI individuals as a function of genomic position, with the gray ribbon indicating standard deviation. The background heatmap displays the number of individuals devoid of sequencing reads as a function of genomic position, with darker shades of red indicating a greater number of individuals with no sequencing reads.

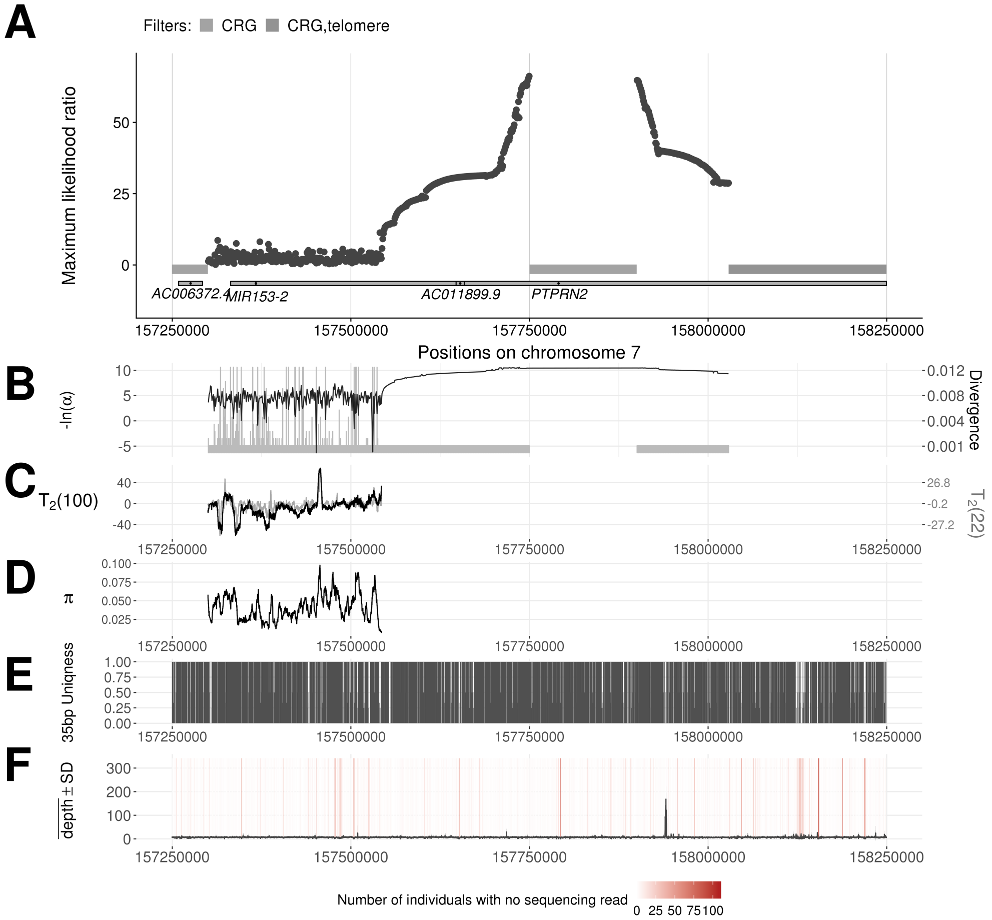

**Fig. S3.6 Introgression sweep signals, parameter estimates, and sequencing properties across the one Mb region on chromosome 19 covering region surrounding *PCAT19* and *CEACAM4* genes in YRI.** **A.** Likelihood ratio test statistic computed from Model 1 of `VolcanoFinder` on data on within-YRI polymorphism and substitutions with respect to chimpanzee. Horizontal dark gray and light gray bars correspond to regions that were filtered based on Hardy-Weinberg equilibrium (HWE) test. Gene tracts and labels for key genes are depicted below the plot, with the wider bars representing exons. **B.** Values for $\alpha$ and divergence $D$ corresponding to the maximum likelihood estimate of the data. Black line corresponds to $-\ln(\alpha)$ and vertical gray bars correspond to estimated $D$. **C.** Likelihood ratio test statistic computed from $T_2$ of `BALLET` on data on within-YRI polymorphism and substitutions with respect to chimpanzee using windows of 100 (black) or 22 (gray) informative sites on either side of the test site. **D.** Mean pairwise sequence difference ($\hat{\theta}_\pi$) computed in five kb windows centered on each polymorphic site. **E.** Mappability uniqueness scores for 35 nucleotide sequences across the region. **F.** Mean sequencing depth across the 108 YRI individuals as a function of genomic position, with the gray ribbon indicating standard deviation. The background heatmap displays the number of individuals devoid of sequencing reads as a function of genomic position, with darker shades of red indicating a greater number of individuals with no sequencing reads.

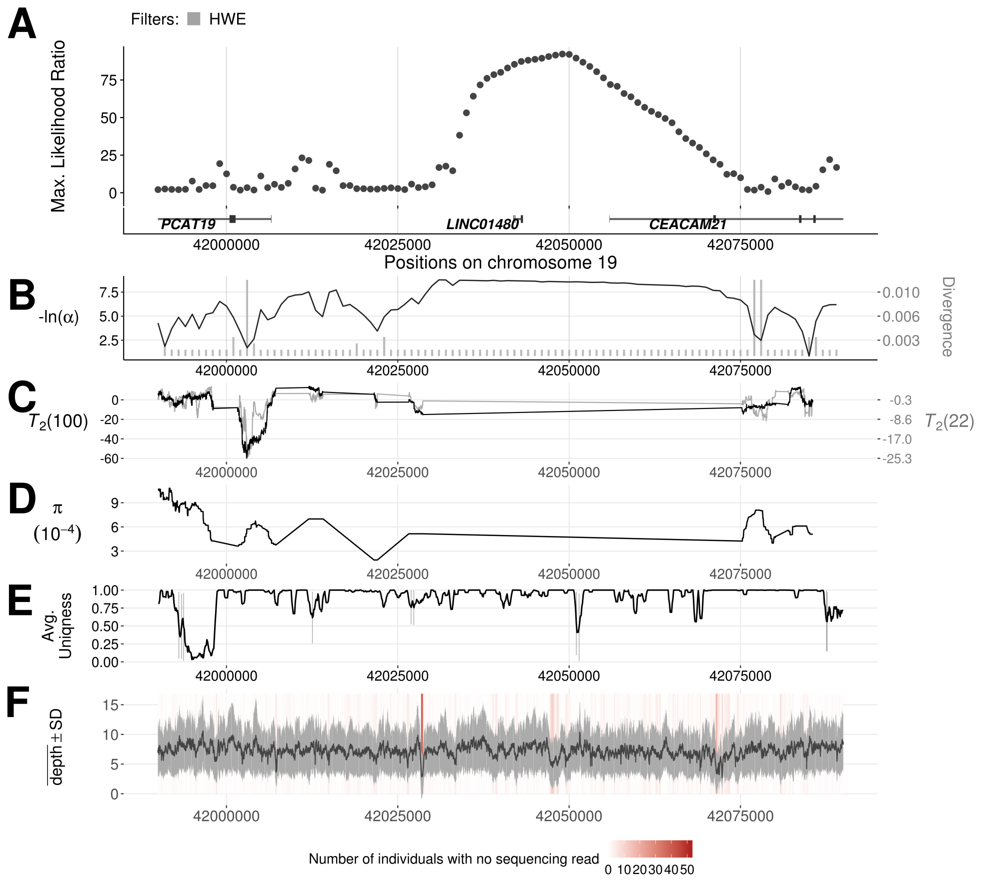

**Fig. S3.7 Introgression sweep signals, parameter estimates, and sequencing properties across the one Mb region on chromosome 17 covering *IGFBP1* and *B4GALNT2* in YRI.**

**A.** Likelihood ratio test statistic computed from Model 1 of `VolcanoFinder` on data on within-YRI polymorphism and substitutions with respect to chimpanzee. Horizontal dark gray and light gray bars correspond to regions that were filtered based on Hardy-Weinberg equilibrium (HWE) test. Gene tracts and labels for key genes are depicted below the plot, with wider bars representing exons. **B.** Values for $\alpha$ and divergence $D$ corresponding to the maximum likelihood estimate of the data. Black line corresponds to $-\ln(\alpha)$ and vertical gray bars correspond to estimated $D$. **C.** Likelihood ratio test statistic computed from $T_2$ of `BALLET` on data on within-YRI polymorphism and substitutions with respect to chimpanzee using windows of 100 (black) or 22 (gray) informative sites on either side of the test site. **D.** Mean pairwise sequence difference ($\hat{\theta}_\pi$) computed in five kb windows centered on each polymorphic site. **E.** Mappability uniqueness scores for 35 nucleotide sequences across the region. **F.** Mean sequencing depth across the 108 YRI individuals as a function of genomic position, with the gray ribbon indicating standard deviation. The background heatmap displays the number of individuals devoid of sequencing reads as a function of genomic position, with darker shades of red indicating a greater number of individuals with no sequencing reads.

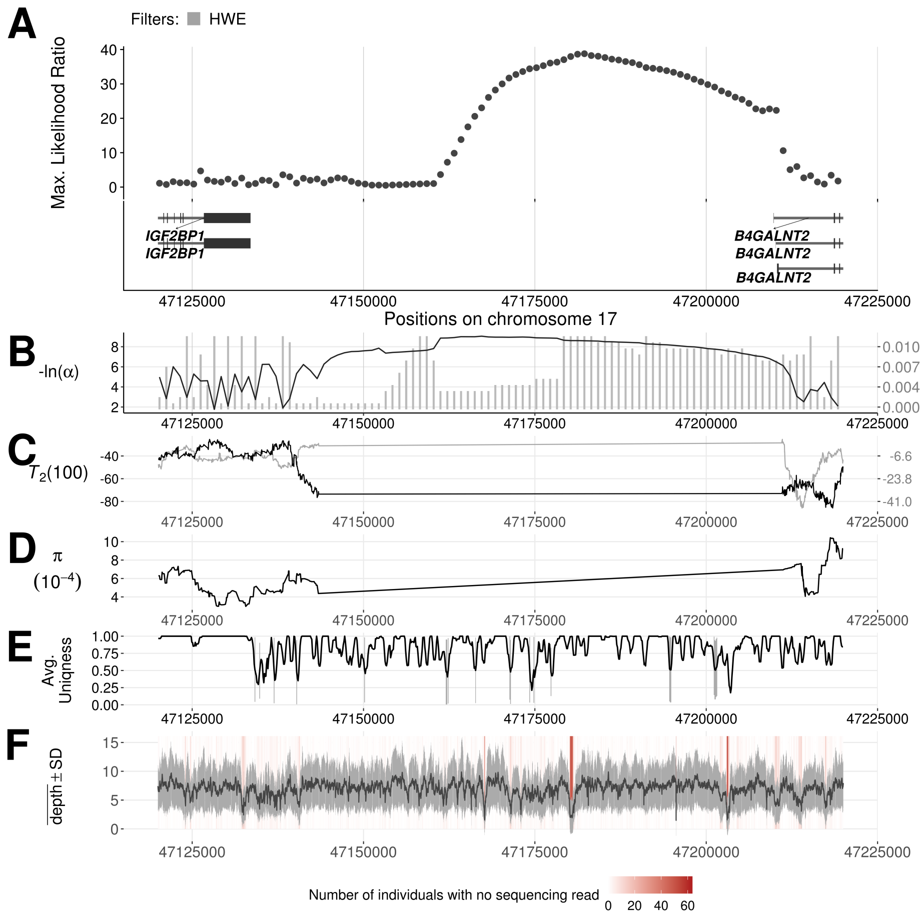

**Fig. S3.8 Introgression sweep signals, parameter estimates, and sequencing properties across the 100 kb region on chromosome 19 covering the gene *MUC4* in CEU.**

**A.** Likelihood ratio statistic computed from Model 1 of `VolcanoFinder` on the data of within-CEU polymorphism and substitutions with respect to the chimpanzee. Gray bars immediately below indicate the type of filters, and the longest gene transcripts are depicted with thick bars standing for exons. **B.** Values for $\alpha$ and divergence $D$ corresponding to the maximum likelihood estimate of the data. Black line corresponds to $-\ln(\alpha)$ and vertical gray bars correspond to estimated $D$. **C.** Likelihood ratio test statistic computed from $T_2$ of `BALLET` on data on within-CEU polymorphism and substitutions with respect to chimpanzee using windows of 100 (black) or 22 (gray) informative sites on either side of the test site. **D.** Mean pairwise sequence difference ($\hat{\theta}_\pi$) computed in five kb windows centered on each polymorphic site. **E.** Mappability uniqueness scores for 35 nucleotide sequences across the region. **F.** Mean sequencing depth across the 99 CEU individuals as a function of genomic position, with the gray ribbon indicating standard deviation. The background heatmap displays the number of individuals devoid of sequencing reads as a function of genomic position, with darker shades of red indicating a greater number of individuals with no sequencing reads.

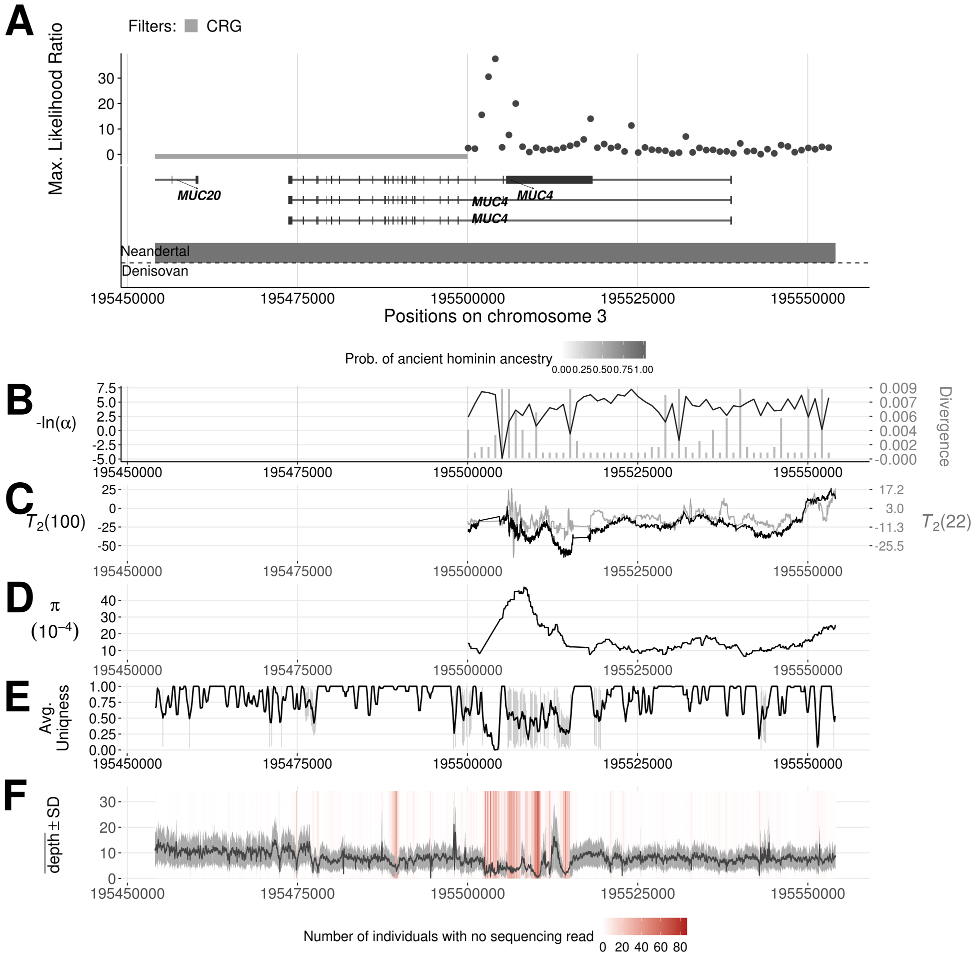

**Fig. S3.9 Introgression sweep signals, parameter estimates, and sequencing properties across the 100 kb region on chromosome 19 covering the gene *CYP2B6* and *CYP2B7* in YRI.**

**A.** Likelihood ratio statistic computed from Model 1 of `VolcanoFinder` on the data of within-YRI polymorphism and substitutions with respect to the chimpanzee. Gray bars immediately below indicate the type of filters, and the longest gene transcripts are depicted with the wider bars standing for exons. **B.** Values for $\alpha$ and divergence $D$ corresponding to the maximum likelihood estimate of the data. Black line corresponds to $-\ln(\alpha)$ and vertical gray bars correspond to estimated $D$. **C.** Likelihood ratio test statistic computed from $T_2$ of `BALLET` on data on within-YRI polymorphism and substitutions with respect to chimpanzee using windows of 100 (black) or 22 (gray) informative sites on either side of the test site. **D.** Mean pairwise sequence difference $(\hat{\theta}_\pi)$ computed in five kb windows centered on each polymorphic site. **E.** Mappability uniqueness scores for 35 nucleotide sequences across the region. **F.** Mean sequencing depth across the 108 YRI individuals as a function of genomic position, with the gray ribbon indicating standard deviation. The background heatmap displays the number of individuals devoid of sequencing reads as a function of genomic position, with darker shades of red indicating a greater number of individuals with no sequencing reads.

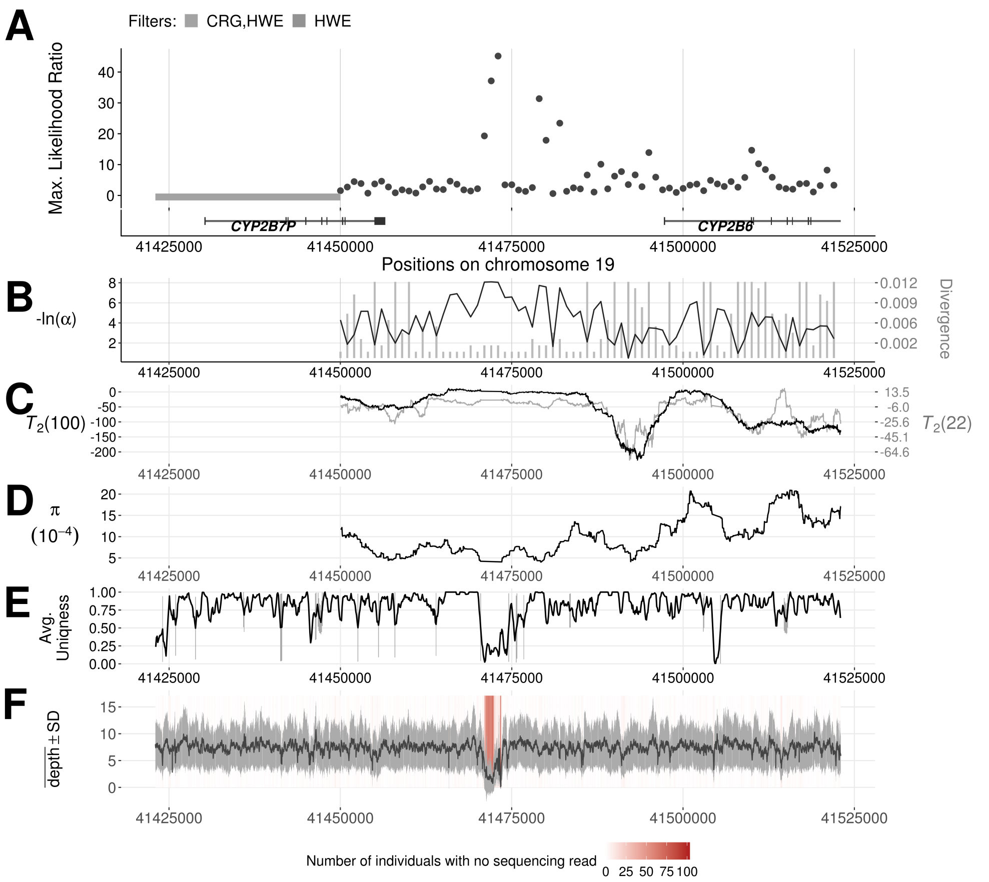
